# Dynamic Mechanochemical feedback between curved membranes and BAR protein self-organization

**DOI:** 10.1101/2020.09.23.310169

**Authors:** Anabel-Lise Le Roux, Caterina Tozzi, Nikhil Walani, Xarxa Quiroga, Dobryna Zalvidea, Xavier Trepat, Margarita Staykova, Marino Arroyo, Pere Roca-Cusachs

**Affiliations:** Institute for Bioengineering of Catalonia (IBEC), the Barcelona Institute of Technology (BIST), 08028 Barcelona, Spain; Universitat Politècnica de Catalunya (UPC), Campus Nord, Carrer de Jordi Girona, 1, 3, 08034 Barcelona; Universitat de Barcelona, 08036 Barcelona, Spain; Institució Catalana de Recerca i Estudis Avançats (ICREA), Passeig de Lluís Companys, 23, 08010 Barcelona; Centro de Investigación Biomédica en Red de Cáncer (CIBERONC), 08028 Barcelona, Spain; Department of Physics, University of Durham, UK; Centre International de Mètodes Numèrics en Enginyeria (CIMNE), 08034 Barcelona, Spain

**Author notes:** These two authors equally participated to the manuscript (Anabel-Lise Le Roux, Caterina Tozzi). **Corresponding authors**, Correspondence to Pere Roca-Cusachs, Marino Arroyo or Anabel-Lise Le Roux.

## Abstract

In many physiological situations, BAR proteins interact with, and reshape, pre-existing curved membranes, contributing to essential cellular processes. However, the non-equilibrium and time-dependent process of reshaping, and its dependence on initial membrane shape, remains largely unknown. Here we explain, both experimentally and through modelling, how a BAR protein dynamically interacts with mechanically bent lipid membranes. We capture protein binding to curved membranes, and characterize a variety of dynamical reshaping events depending on membrane shape and protein arrangement. The events can be generally understood by an isotropic-to-nematic phase transition, in which low curvature templates with isotropic protein orientation progress towards highly curved lipid tubes with nematic protein arrangement. Our findings also apply in cells, where mechanical stretch triggers BAR-protein-membrane interactions that enable potential mechanotransduction mechanisms. Our results characterize and broaden the reshaping processes of BAR proteins on mechanically constrained membranes, demonstrating the interplay between membrane mechanical stimuli and BAR protein response.

---

Due to the curved shape and membrane binding of Bin/Amphiphysin/Rvs (BAR) domains, proteins containing such domains have the interesting ability to reshape membranes. Furthermore, because of their elongated shape, they can align along a preferred direction, adopting a nematic organization that impinges anisotropic curvature on the membrane. For instance, incubation of small vesicles with a high concentration of BAR proteins leads to tubules covered by a dense protein scaffold where the elongated molecules are nematically arranged^1,2^. In a different system, GUVs with sufficiently high bound protein density rapidly expel thin protein-rich tubes in a tension dependent manner^3^. On thin membrane tubes pulled out of giant unilamellar vesicles (GUVs), BAR proteins can also change the radius of the tube and the force required to hold it^4^. Beyond these well-known cases, in many physiological situations BAR proteins interact with pre-existing curved membrane templates. Such templates can include for instance invaginations caused by nanoscale topographical features on the cell substrate^5^, mechanical folds^6,7^, or endocytic structures^8,9^. Due to their affinity for curved membranes, BAR proteins are thus bound to reshape such templates in ways that could be important in processes ranging from endocytosis to the sensing of topographical or mechanical cues in the cell environment. However, the dynamics of membrane reshaping by BAR proteins, and how it depends on initial membrane shape, remains elusive.

To address this issue, we developed a novel experimental system combined with theoretical and computational modeling to study the reshaping of cellular-like membrane structures of a broad range of shapes and sizes. In our system, we create curved membrane features off a supported lipid bilayer by lateral mechanical compression. As previously shown in vitro^6^ and in cells^7^, once stretch is applied to membranes and subsequently released, excess membrane area is stored in protrusions of tubular or spherical shape. The size and shape of tubes or spherical caps depends on the relative magnitude of excess area and excess enclosed volume. Controlling stretch, but also osmolarity, it is thus possible to generate tubules, buds, or spherical caps. In contrast with tubes pulled out of GUVs, where a tip force and tension are required to stabilize their shape, in our system tubes are stabilized osmotically without a pulling force. These protrusions emerging from a flat supported bilayer can serve as model system for membrane templates such as endocytic buds, or topographically/mechanically induced structures.

Experimentally, we used the liposome deposition method to form a fluorescently labelled supported lipid bilayer (SLB) on top of a thin extensible polydimethylsiloxane (PDMS) membrane. To this end, an electron microscopy grid was deposited on top of the PDMS membrane before plasma cleaning, which activated only the uncovered PDMS areas^10^. An easily identifiable hexagonal pattern was obtained (Fig. 1a), with a fluid SLB formed inside the hexagon, (Suppl. Fig. 1) while a lipid monolayer was formed outside. The membrane was placed inside a stretching device previously described^7^ and mounted on a spinning disk confocal microscope (Fig. 1a). At initial state, the fluid bilayer contained brighter signals coming from non-fused liposomes (Suppl. Fig. 2a). The patterned SLB (pSLB) was then uniformly and isotropically stretched for 120 s (until 5 to 8 % strain), slowly enough to allow liposome incorporation in the strained fluid bilayer, thereby ensuring membrane integrity (as happens in a cellular membrane through lipid reserve incorporation^7^). After 120 s, stretch was slowly released during 300 s to a completely relaxed state, and lateral compression led to the formation of lipid tubes and buds (Fig. 1b, Suppl. Fig. 2a and Suppl. Videos 1 and 2). We note that our system is diffraction-limited and not amenable to electron microscopy, and we could thus not measure tube diameter.

**Figure 1:**
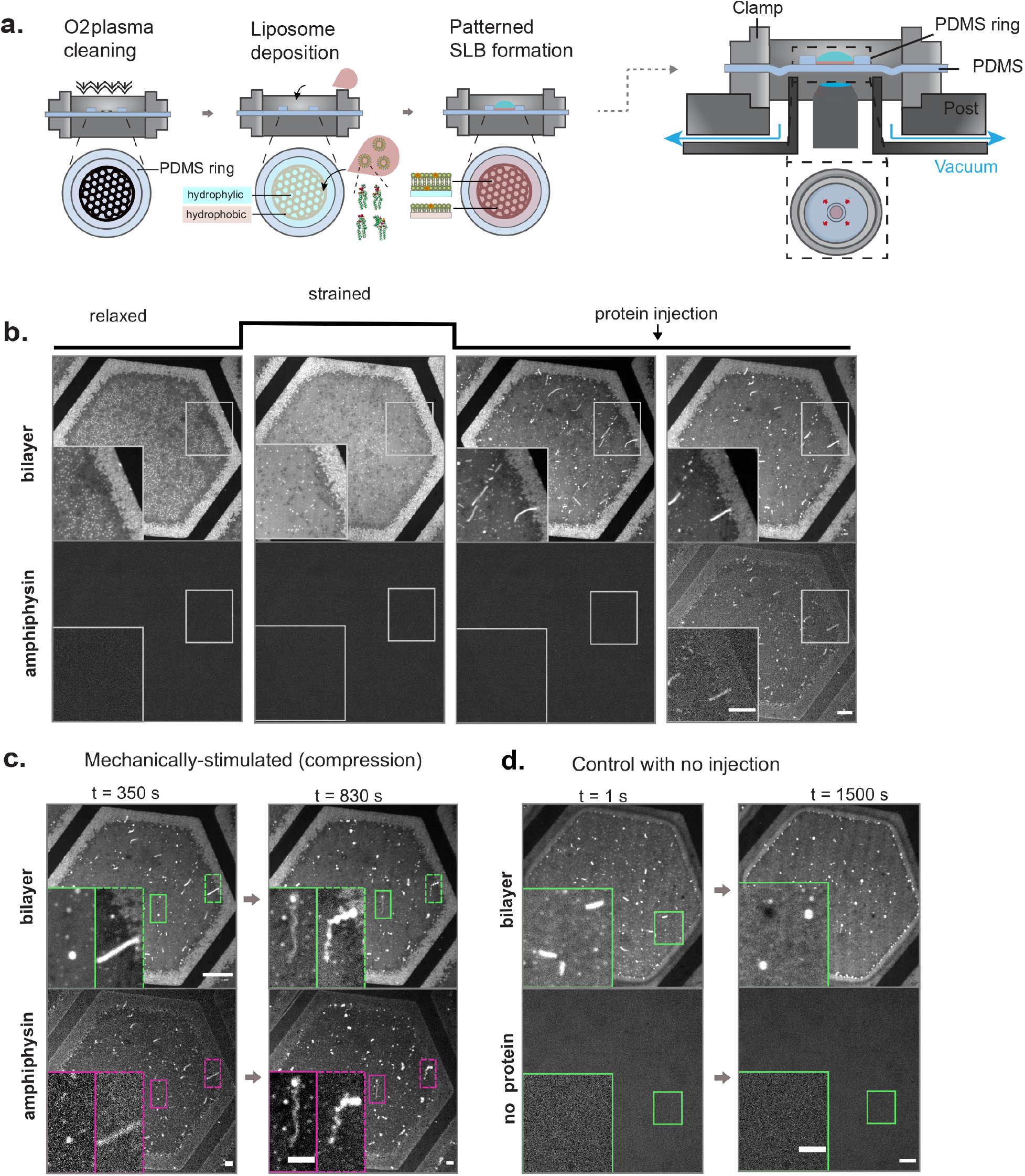
Experimental system. **a**, Schematics of the patterned supported lipid bilayer (pSLB) placed in a stretch system compatible with confocal microscopy. The pSLB is obtained by plasma cleaning a PDMS membrane in presence of a TEM grid. Only the exposed PDMS becomes hydrophilic, and subsequent liposome deposition renders a SLB after buffer rinse. The non-exposed PDMS remains hydrophobic and a lipid monolayer is formed instead. **b**, Representative images of the mechanical stimulation of the pSLB, showing both lipid and protein fluorescence images. In the resting initial state, excess liposomes stand on top of the pSLB. With strain, the liposomes incorporate in the pSLB. Upon release, excess lipids are expelled in the form or tubes or buds. At this stage, fluorescent Amphiphysin is gently microinjected on top of pSLB and its binding to the tubes and buds is monitored with time. **c**, Membrane tubes (green inset) and buds (purple inset) before (left) and after (right) being reshaped by Amphiphysin. **d**, Control in which no protein is injected on top of the pSLB. Scale bar, 5 *μ*m.

Then, we injected fluorescently labelled Amphiphysin in the bulk solution on top of the pSLB (Fig. 1b). Amphiphysin, an N-BAR protein binding lipid bilayers of positive curvature (invaginations), has been often used as a model for BAR proteins^4,11,12,13,14^, and is known to bind to negatively charged lipids^15^ and especially to 1,2 dioleoyl-sn-glycero-3-phosphate (DOPA)^16^, which may enhance electrostatic interactions. We assessed protein activity by measuring the diameter of Amphiphysin-reshaped tubes via transmission electron microscopy in the sucrose loading vesicle assay^17^. Consistent with the literature, we found diameters of ~25 nm, Suppl. Fig. 2b. The fluorescence signal from both the pSLB and Amphiphysin (in different channels) was subsequently monitored. Once injected, the protein bound to the tubes and buds (Fig. 1c, left) and, after further adsorption from the bulk, it started to reshape them into geometrically heterogeneous structures with coexistence of small spherical and tubular features (Fig. 1c, right). As controls, we first monitored the tubes in absence of protein injection; they were stable for some minutes and then started to relax to buds standing on top of the bilayer, presumably due to enclosed fluid and/or excess membrane reorganization within the system (Fig. 1d and Suppl. Video 3). Additionally, we performed the same experiment by injecting fluorescent neutravidin instead of Amphiphysin. Neutravidin did not specifically bind to the tube, and no reshaping was observed apart from the same tube-to-bud relaxation observed in the absence of injection (Suppl. Fig. 2c and Suppl. Video 4). Finally, we monitored the effect of Amphiphysin in non-stretched membranes, which were therefore devoid of pre-existing membrane structures. In this case, membrane reshaping only occurred if Amphiphysin concentration was increased above 5 μM in the bulk, which merely consisted in the formation of bright/dark spots in the membrane, likely reflecting membrane tearing. (Suppl. Fig. 2d, e and Suppl. Videos 5 and 6).

To understand the physical mechanisms underlying our observations, we developed a theoretical framework considering the dynamics of lipid tubes and buds with low coverage (since protein is injected once structures are formed) and low curvature (since the structures are made markedly thinner by Amphiphysin) upon exposure to BAR proteins. Theoretically, various computational studies using coarse-grained simulations of elongated and curved objects moving on a deformable membrane have suggested the self-organization of regions with high anisotropic (cylindrical) curvature with high-protein coverage and strong nematic order ^18,19,20^. None of these works, however, predicted or observed the tube-sphere complexes that appear in our experiments (Fig. 1c). To address this, we first developed a mean field density functional theory for the free energy of the proteins *F_prot_*. This theory accounts for protein area coverage *ϕ*, orientational order as given by a nematic order parameter *S*, and membrane curvature. This is presented in detail in an accompanying paper^21^.

On flat membranes and for elliptical particles of the size and aspect ratio of Amphiphysin, the theory predicts an entropically-controlled discontinuous isotropic-to-nematic transition during which the system abruptly changes from low to high order as protein coverage increases above *ϕ* ≈ 0.5, in agreement with previous results in 3D^22^, Suppl. Fig. S3a1. On curved surfaces, our model also accounts for the elastic curvature energy of proteins, which depends on the curvature of the surface, and on their intrinsic curvature and orientation relative to the surface directions of principal curvatures (Fig. 2a). We then examined the protein free-energy landscape on spherical surfaces, which according to the theory coincides with that of the flat membrane with a bias proportional to *ϕ* times the bending energy of proteins on the curved surface. Thus, the minimum energy paths as density increases (red dots in Fig. 2b-e) and hence the abrupt isotropic-to-nematic transition persist regardless of sphere radius, Fig. 2b, c, noting that on a complete sphere the nematic phase necessarily involves defects^18^. On cylindrical surfaces, however, curvature is anisotropic and the energy landscape is fundamentally modified according to our theory as proteins can lower their free energy by orienting along a direction of favorable curvature. The competition between protein bending and entropy results in a continuous isotropic-to-nematic transition (Fig. 2d) and a significant degree of orientational order even at low coverage when the tube curvature is comparable to that of the protein (Fig. 2e). The model thus predicts how the nematic ordering of the curved and elongated membrane depends on coverage, curvature, and curvature anisotropy.

**Figure 2:**
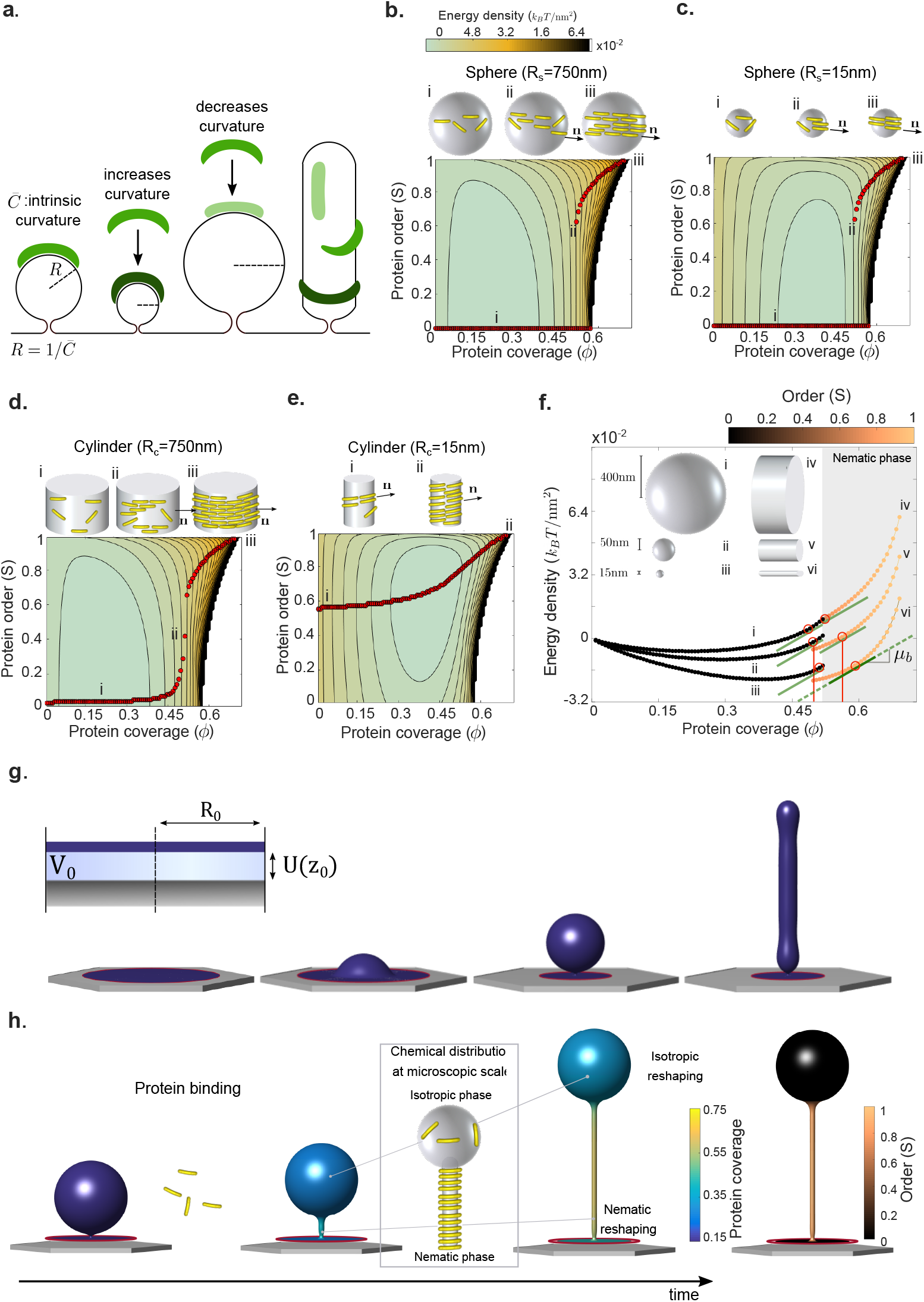
Theoretical and computational modeling. **a**, Schematic diagram of a BAR domain interacting with a lipid membrane. Protein elastic energy depends on surface curvature and protein orientation. For cylindrical surface, curvature is maximal (dark green) and minimal (light green) along perpendicular directions. **b-e**, Energy density landscape according to our mean field density functional theory depending on protein coverage *ϕ*, nematic alignment *S*, and the shape and size of the underlying membrane (sphere or cylinder as illustrated on top of each plot). Red dots denote states of equilibrium alignments S for a given protein coverage ϕ, i.e. minimizers of the free energy along vertical profiles, depicting the transition from isotropic (i) to nematic phase (ii-iii). The white region in the energy landscape is forbidden due to crowding. **b-c**, show discontinuous transitions for protein alignment on isotropically curved membranes. **d-e**, show continuous transitions for anisotropically curved membrane. The intrinsic protein radius of curvature is 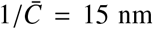. **f**, Free energy profiles for spheres and cylinders of different sizes along the equilibrium paths. The chemical potential of proteins is the slope of these curves. All points marked with red circles have the same chemical potential at the tangent points *μ_b_* and hence are in chemical equilibrium. **g**, Membrane protrusions obtained by lateral compression of an adhered membrane patch of radius *R*_0_ interacting with a substrate with a potential *U*(*z*) and for various amounts of enclosed volume *V*_0_, see Theoretical Model. **h**, Schematic of reshaping dynamics involving membrane relaxation, and protein binding, diffusion and ordering.

We then studied whether the model predicted the experimentally observed coexistence of thin tubes (which according to the theory should have higher coverage and order) and larger spheres (which should have lower coverage and isotropic organization). We examined the energy landscape along the minimizing paths (red dots) for spheres and tubes of varying radius (Fig. 2f). Since the slope of these curves is the chemical potential of proteins on the membrane, which tends to equilibrate with the fixed chemical potential of dissolved proteins in the medium, points of chemical coexistence are characterized by a common slope (red circles). This figure shows the largely non-unique combinations of geometry and membrane coverage compatible with coexistence in chemical equilibrium between higher-coverage nematic phases on cylinders and lower-coverage isotropic phases on spheres, supporting plausibility of such coexistence in the dynamical structures.

Shape, however, is also a dynamical variable and the selection of coverage and shape requires the two-way interplay between the chemical free energy *F_prot_* and the elastic free energy of the membrane *F_mem_.* To account for this and for the out-of-equilibrium nature of our experiments, we took the mean field energy density functional theory, as a foundation to develop a self-consistent continuum chemo-mechanical model. This model predicts the interplay between protein behavior and membrane shape, and accounts for *F_prot_* + *F_mem_*, for the dynamics of protein adsorption from a bulk reservoir, for the diffusion of proteins on the surface, and for the membrane dissipation associated with shape changes^23^ (see Methods, Theoretical Model, for a discussion of the model and its parameters). Starting from membrane protrusions in mechanical equilibrium (tubular or spherical, Fig. 2g) off a supported bilayer in the absence of proteins and for a fixed membrane tension and enclosed volume^6,7^, this model predicts the dynamics of membrane shape, *ϕ*, and *S* following a sudden increase of dissolved protein concentration in the medium (Fig. 2h).

We then tested different model predictions using the experimental setup. First, we considered the dynamics of a spherical bud connected to the supported bilayer by a neck. As per the model, proteins should adsorb on the entire membrane but at a faster rate at the neck, where membrane curvature is more favorable than on the vesicles or on the flat part. Furthermore, due to a local gradient in chemical potential, proteins are further recruited by diffusion towards the neck, where they rapidly adopt a nematic order (Fig. 2e) in contrast with the isotropic order in the vesicle. The lower energy of proteins on the thin neck (Fig. 2f) outweighs both the higher membrane curvature energy of a tube relative to a larger vesicle and the entropic penalty of a local protein enrichment. This leads to a progressive elongation of the neck into a thin tube with higher coverage and nematic order (Fig. 2h, 3a and Suppl. Video 7). If the protrusion is allowed to exchange enclosed volume with the adhered part of the membrane (see Methods, Theoretical Model), tube elongation occurs at the expense of the vesicle area (Suppl. Fig. 4a and Suppl. Video 8). According to our simulations, the radius of these thin tubes is of about 15 nm, close to the radius of tubes scaffolded by Amphiphysin (Suppl. Fig. 2b and 4b). Consistent with model predictions, the experimental observations systematically captured the growth of thin necks connecting shrinking vesicles to the supported bilayer (Fig. 3a, Suppl. Fig. 4c and Suppl. Video 9). The contrast of nematic order between vesicles and tubes predicted by our simulations was not accessible experimentally. However, another hallmark of the isotropic-nematic coexistence suggested by the model is a protein enrichment on the tube relative to the vesicle. For a wide range of bud diameters and protein concentrations, our simulations predicted a stable and approximately two-fold higher protein concentration in tubular versus bud regions (Fig. 3b and Suppl. Fig. 5b). When we estimated this experimentally (Suppl. Fig. 5a), this enrichment was confirmed (Fig. 3b).

**Figure 3:**
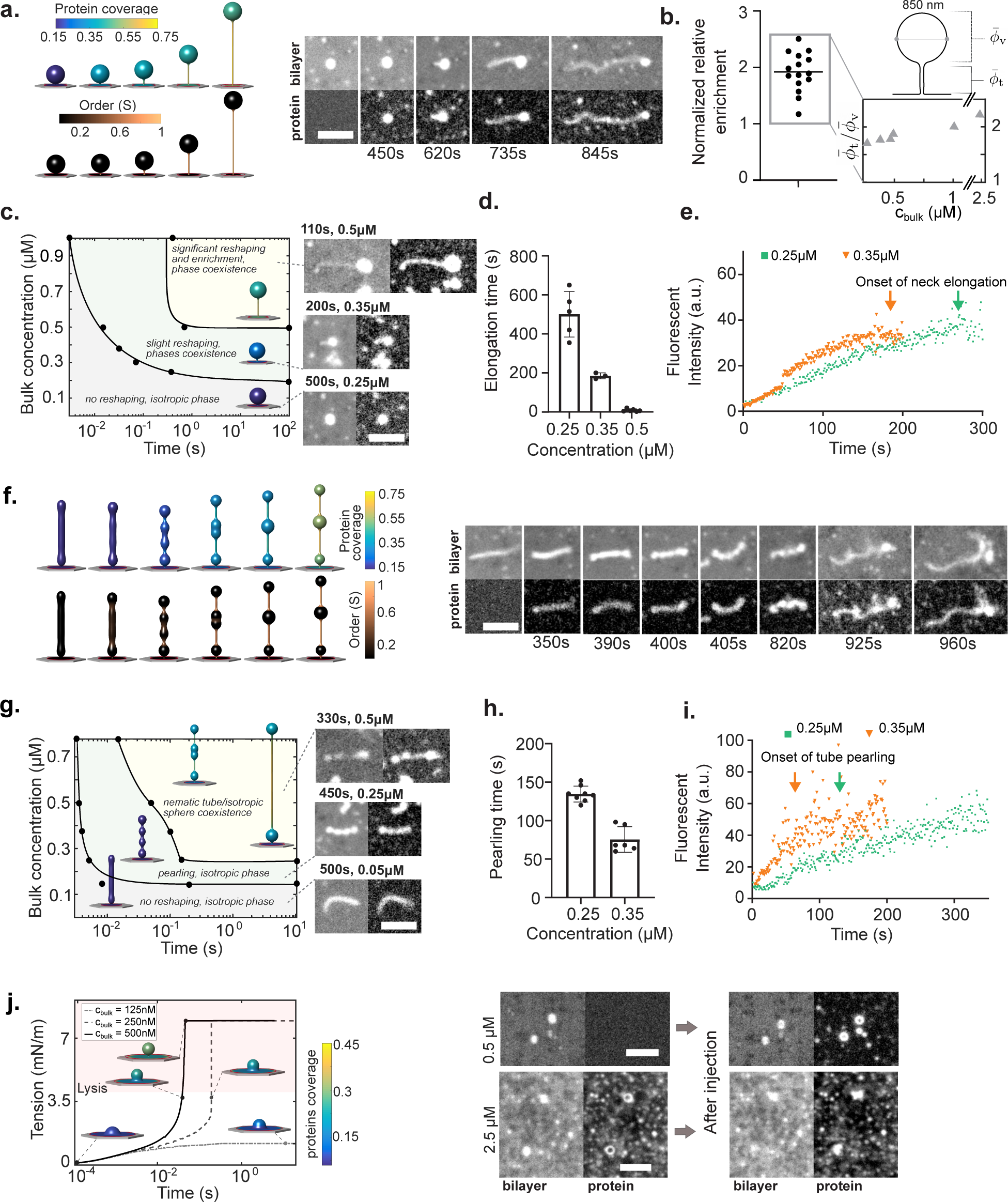
Dynamics of membrane reshaping. **a**, Results of simulations (left) and experiments (right) showing bud reshaping with time in response to Amphiphysin. **b**, Ratios of protein coverage on tubes versus buds, normalized to the values measured for the lipid bilayer. Left, experimental values (n=15), right, theoretical concentration ratios *ϕ_t_/ϕ_v_* for a 0.85 *μ*m diameter bud, exposed to different bulk concentrations. **c**, Dynamical diagram of bud reshaping as a function of time and bulk protein concentration, classifying the state of the system as one with no reshaping and isotropic protein organization (*S* ≈ 0), one with slight tube elongation, pearling, some degree of order (*S* < 0.5) and low coverage, and one with significant elongation, enrichment and phase coexistence. Inserts are experimental examples. **d**, Times at which bud elongation starts as a function of concentration (n=5,3 and 6 for 0.25, 0.35 and 0.5 *μ*M respectively). **e**, Examples of Amphiphysin fluorescence intensities in buds incubated at two different concentrations. Bud elongation times are marked with an arrow. **f**, Results of simulations (left) and experiments (right) showing tube reshaping with time in response to Amphiphysin. **g**, Dynamical diagram of tube reshaping as a function of time and bulk protein concentration. **h**, Times at which tube pearling starts as a function of concentration (n=5 and 6 for 0.25 and 0.35 *μ*M respectively). **i**, Examples of Amphiphysin fluorescence intensities in tubes incubated at two different concentrations. Tube pearling times are marked with an arrow. **j**, Left, Model prediction in pressurized caps of about 400 nm in radius exposed to different Amphiphysin concentrations. States in the pink shaded area are prone to membrane lysis. Right, Initial and final states of pressurized caps (obtained from an hypoosmotic shock) upon incubation with Amphiphysin. At 2.5 *μ*M concentration, lysis of the caps can be observed. Scale bars, 5 *μ*m.

Experimentally, we found that a threshold bulk concentration was required for the growth of such tubes within the observed time frame (no growth observed at 0.05 μM after 1100s, Suppl. Fig. 4c). We also found that the higher the concentration, the faster reshaping occurred. We thus systematically studied how increasing concentrations of protein in the bulk affected model outcome and reshaping over time. This led to a dynamical diagram of bud shape as a function of time and protein concentration (Fig. 3c and Suppl. Fig. 4b). We note that we lacked a precise experimental control of the dynamics of protein delivery to the membrane due to diffusive and possibly advective transport from the injection to the observation point, and hence a fully quantitative comparison between theory and experiment regarding the time-scales of reshaping was not possible. However, our simulations assuming instantaneous exposure of protein in solution also show a strong concentration dependence in the reshaping dynamics (Fig. 3c). Increasing protein concentration decreases the time required to initiate tubulation, representative of the time at which proteins nematically arrange. This was experimentally confirmed by measuring the times of tubulation initiation (Fig. 3d) and by looking at the protein binding curves on the buds. Such curves were obtained by plotting the mean intensity of the protein fluorescence on buds over time, until tubulation starts. The protein concentration triggering tubulation was reached faster at higher concentration (Fig. 3e and compare Suppl. Videos 9 and 10, in which tube elongation starts much faster at 0.5 μM bulk Amphiphysin concentration versus 0.25 μM). Representative images of the reshaped state of buds at increasing times are plotted in Suppl. Fig. 4c. As an additional control, we noted that bud elongation also occurred upon exposure to non-fluorescently labelled Amphiphysin (Suppl. Fig. 4d).

Then, we considered the reshaping dynamics of tubes, which were frequently formed upon compression. For low protein concentration, proteins adsorbed onto the tubes and did not lead to an observable reshaping. Yet, the protein had a stabilizing effect on tubes since, unlike those in the absence of proteins, most of them did not spontaneously relax into buds (as shown in Suppl. Fig. 6c where the tube exposed to 0.05 μM bulk protein concentration was still stable after 1100s.). At higher concentrations, however, we systematically found that tube reshaping was initiated by the formation of a sequence of pearls (Fig. 3f and Suppl. Video 11 and 12). Previous results have shown the formation of pearled membrane tubes as a result of a sufficiently large isotropic and uniform spontaneous curvature^24,23,25^. Indeed, we hypothesized and our simulations show that, if initial tubes were large-enough, then at low coverage they should exhibit a largely uniform and isotropic arrangement of molecules, Fig. 2d, e, hence impinging an isotropic spontaneous curvature on the membrane leading to pearling (Suppl. Fig. 6a; in the absence of a nematic transition, no further reshaping would follow this pearling phase). The pearling instability produces several membrane necks along the tube. If coverage is high enough, sufficiently many proteins may be drawn to those necks. This triggers an isotropic-to-nematic transition, a progressive elongation of thin tubes, and a consumption of spheres analogous to that described above in membrane buds (Fig. 3f).

Analogously to the case of bud elongation, we built a dynamical diagram of tube shape as a function of bulk protein concentration and time (Fig. 3g and Suppl. Fig. 6b). Increasing concentration decreases the time to initiate pearling, which was also observed experimentally (Fig. 3h, i and compare the earlier tube pearling observed in Suppl. Video 12, 0.25 μM bulk Amphiphysin concentration, with Suppl. Video 13 at 0.35 μM.). Concentration also accelerates the subsequent transitions from uniform to pearled tubes, then to pearls connected by tubes (Fig. 3g and Suppl. Fig. 6b, c). This configuration was stable for long times in simulations and experiments (though such reshaped tubes collapsed on themselves, likely due to the related loss of tension^3,26^ — this phase is best observed in the movie, see Suppl. Video 13). As in the case of bud reshaping, tube pearling also occurred upon exposure to non-fluorescently labelled Amphiphysin (Suppl. Fig. 6d).

For both tubes and buds, we evaluated the protein coverage required to trigger reshaping. Though measuring protein coverage in our experimental set up is very challenging, we developed a protocol to obtain an estimate. We performed a classical calibration of the protein fluorescence versus coverage^27^, and subsequently corrected the data by a geometrical factor taking into account the out-of-plane loss of signal and the geometrical signal integration of non-planar structures (Suppl. Fig. 7a). As a result, initiation of bud elongation or tube pearling occurred at ~0.4 and ~0.25-0.35 coverage respectively, approximately matching theoretical predictions (Suppl. Fig. 7b). Taken together, our results show that membrane curved templates exposed to sufficiently high concentrations of BAR proteins evolve towards uniformly thin and protein-rich nematic tubes. During the process, heterogeneous intermediates are formed, exhibiting mixtures of low curvature and low isotropic coverage regions with others of high cylindrical curvature, high coverage, and nematic order.

Finally, we evaluated the case of shallow spherical cap protrusions, which develop when hypoosmotic shocks are generated both in vitro and in cells^6,7^. In this case, the membrane needs to accommodate a significant excess volume of liquid with little excess membrane area, leading to a structure under significant tension^6^. Under these conditions of small excess area, shape changes are very difficult. Our model predicts that upon exposure of such shallow caps to BAR proteins, shape changes are negligible. Instead, tension in the membrane sharply increases, potentially leading to membrane tearing^28,12^ (Fig. 3j). Accordingly (acknowledging as before that direct comparison of concentrations in experiments and simulations is not straightforward), shallow spherical caps formed by a hypo-osmotic shock in our experimental system were not visibly reshaped by Amphiphysin even at significant concentrations (Fig. 3j and Suppl. Video 14), and teared and collapsed upon exposure to higher Amphiphysin concentrations (Fig. 3j and Suppl. Video 15).

Beyond the specifics of the reshaping process, an important conclusion from this study is that the mechanical generation of membrane structures acts as a catalyzer of membrane reshaping by BAR-domain proteins. Indeed, compressed membranes exhibited a wide range of reshaping behaviors (Fig. 3), whereas non-mechanically stimulated membranes exposed to the same Amphiphysin concentration did not reshape in any clear way (Suppl. Fig. 2d and Suppl. Video 5). This suggests the interesting possibility that cells could harness the mechanically-induced formation of membrane invaginations^7^ to trigger BAR-mediated responses, thereby enabling mechanosensing mechanisms. To explore this possibility, we cultured dermal fibroblasts (DF) and overexpressed GFP-Amphiphysin, which is well known to trigger spontaneous membrane tabulation^11^. Then, we stretched and subsequently compressed the cells using a previously described protocol^7^. Upon compression, cells formed dot-like membrane folds termed “reservoirs” (Fig. 4a), analogous to the membrane structures observed in vitro in Fig. 1 and Fig. 3. Amphiphysin-containing membrane tubes formed before, during, and after stretch. However, their number decreased during the stretch phase, likely due to increased membrane tension (Fig. 4c). Upon release of the stretch, tube formation strongly increased, reaching values well above the initial non-stretched condition (Fig. 4c and Suppl. Video 16). Further, tubes formed upon destretch nucleated close to reservoir locations (Fig 4b). We measured the elongation rates of these tubes and they ranged from 200 to 350 nm/s, comparable with the elongation rates found in-vitro for bud elongation (from 40 to 600 nm/s, with higher elongation rate at higher concentration). Though Amphiphysin overexpression presumably leads to concentrations above physiological levels, these results clearly show that mechanical compression of cells can stimulate BAR-mediated membrane tubulation.

**Figure 4:**
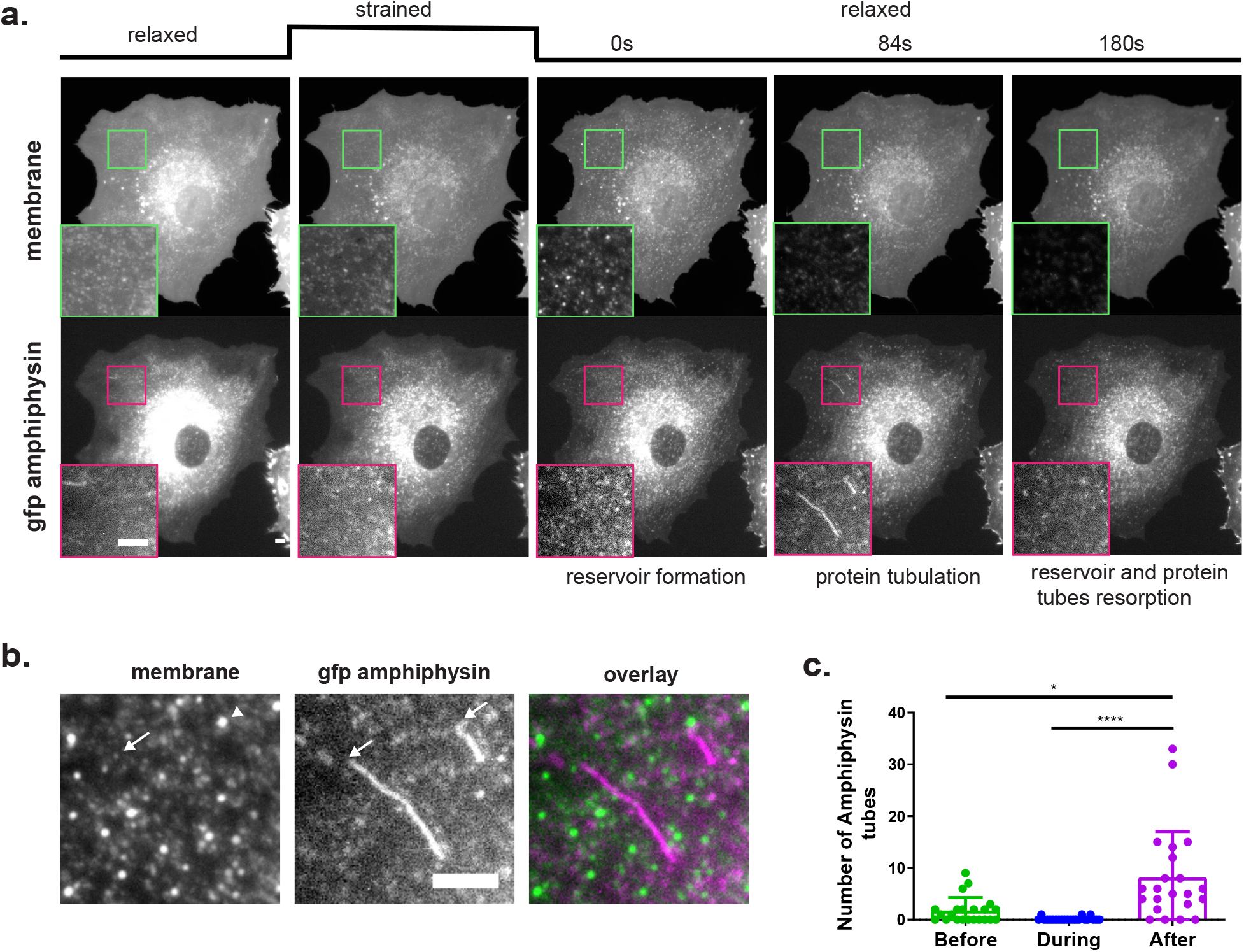
Mechanical stretch in cells triggers Amphiphysin-mediated tubulation. **a**, Representative images (in both membrane and Amphiphysin channels) of a cell before, during and after stretch release. **b**, Detail of membrane and Amphiphysin channels during tubulation. **c**, Quantification of the number of Amphiphysin tubes at rest, during stretch and once stretch is released (n=22,*: p< 0.05, ****: p< 0.0001, Friedman test). Scale bars, 5 *μ*m.

That BAR-domain proteins can reshape membranes is well known, but the dynamics of the process and its dependence on the initial template were unexplored. Here we show that the dynamics of reshaping conforms a very rich process with many intermediate steps, including phase separation between isotropic and nematic phases, and with major reshaping processes occurring at low coverage and curvature. The curvature sensing and membrane reshaping properties of BAR proteins have been extensively studied on highly curved tubes (up to 100 nm in diameter), mostly in equilibrium^4,29,30,31^. However, many cell studies pointed out the role of BAR proteins acting on lower curvature lipid structures^32^. Our study demonstrates and characterizes reshaping at low concentration, low curvature, and low tension, a highly relevant scenario in cells. This behavior emerges naturally from the fundamental physics of membrane mechanics and its mechanochemical interactions with curved proteins, generating a non-trivial feedback between membrane mechanical stimulation and subsequent response. Beyond the physics of the process, such feedback could potentially be used in the many cellular processes involving membrane reshaping under mechanical constraints. This includes the well-studied role of BAR proteins in endocytosis, but also emerging roles in maintenance of cell polarity^33^, response to osmotic changes^34^ or build-up of caveolar structures^35,36^.

## Methods

### Protein expression and purification

The plasmid containing full length human Amphiphysin 1 (FL-hAMPH), pGEX-Amphiphysin1, was a kind gift from Pr. De Camilli, Yale University. The plasmid codes for the FL-hAMPH preceded by a Glutathione S-Transferase (GST-Tag) and a cleavage site recognised by prescission protein. The plasmid was transformed in *Escherichia coli* Rosetta™ (DE3) pLysS cells (Novagen). Selected colonies were grown in luria broth supplemented with 25 μg/ml chloramphenicol and 25 μg/ml kanamycin at 37 C until an OD between 0.6-0.8 was reached. Protein expression was induced by 1 mM isopropyl-β-D-thiogalactopyranosid (IPTG) overnight at 25 C. Cells were pelleted for 30 min at 4000 rpm, pellet was resuspended in lysis buffer (10 mM phosphate buffer saline pH 7.3, supplemented with cOmplete protease inihibitor, EDTA free (Roche) and 1 mM Phenylmethylsulfonyl fluoride (Sigma)). Cells were lysed (5 pulses of 30 s sonication with 30 s rest), incubated for 20 min on ice with 5 μg/ml DNase, and centrifuged at 25000 rpm for 45 min. The supernatant was collected and incubated with 2 column volume (for 20 mL supernatant) of Gluthatione Sepharose 4B (GE Healthcare) for 1h30 on a rotating wheel. The beads were subsequently washed with phosphate buffer saline buffer, pH 7.3 before exchanging to the cleavage buffer (50 mM Tris-base, 150 mM NaCl, 1 mM EDTA, 1 mM DTT, pH 7.0). 60 units of Prescision protease (BioRad Laboratories) were added to the beads and cleavage of the GST-Tag was allowed for 1 h at room temperature followed by an overnight incubation at 4 C on a rotating wheel. The flow through was recovered, and contained cleaved amphiphysin that was further purified by size exclusion chromatography in a Superdex 75 26/60 in 10 mM PBS, pH 7.5, 1 mM DTT. Two fractions were obtained, both containing amphiphysin according to the SDS-page gel, but the second fraction of smaller size was taken and concentrated for further use. The purity and identity of the product was established by HPLC and mass spectrometry (BioSuite pPhenyl 1000RPC 2.0 x 75 mm coupled to a LCT-Premier Waters from GE Healthcare). Neutravidin was from Thermofisher. Proteins (amphiphysin and neutravidin) were coupled to an Alexa Fluor® 488 TFP ester according to the manufacturer protocol and the resulting protein-alexa 488 was concentrated again. Adsorption was measured in a Nanodrop at 280 nm to obtain protein concentration and at 488 nm to obtain fluorophore concentration. This gave an average amount of fluorophore per protein of 3 per amphiphysin dimers and 1 per neutravidin protein. Amphiphysin was frozen and kept at −80 C, experiments were performed with freshly unfrozen samples. Protein integrity was verified by SDS-page of the unfrozen samples.

### Preparation of stretchable membranes

Stretchable polydimethylsiloxane (Sylgard Silicone Elastomer Kit, Dow Corning) membranes were prepared as previously described^7^. Briefly, a mix of 10:1 base to crosslinker ratio was spun for 1 minute at 500 rpm and cured at 65° C overnight on plastic supports. Once polymerized, membranes were peeled off and assembled onto a metal ring that can subsequently be assembled in the stretch device.

### Patterned Supported Lipid Bilayer (pSLB) formation on PDMS membrane

pSLBs were prepared by combining 1,2-dioleoyl-sn-glycero-3-phosphocoline (DOPC), 1,2-dioleoyl-snglycero-3-phospho(1’-rac-glycerol) (sodium salt) (DOPS), and 1,2-dioleoyl-sn-glycero-3-phosphate (sodium salt1,2-dipalmitoyl-sn-glycero-3-phosphoethanolamine-N-(lissamine rhodamine B sulfonyl) (ammonium salt) (LissRhod-DPPE). 1.25 mg of total lipids in a DOPC:DOPS:DOPA 3:2:1 proportion, with 0.5 % mol LissRhod-DPPE were dissolved in chloroform. The solvent was evaporated for minimum 4 h. The lipid film was immediately hydrated with 750 μL of PBS, pH 7.5 (final concentration of 1.6 mg/ml) at room temperature. After gentle vortexing, a solution of giant multilamellar vesicles was obtained. Large unilamellar vesicles (LUVs) were prepared by mechanical extrusion using the Avanti extruder set. The lipid suspension was extruded repeatedly (15 times) through a polycarbonate membrane (Whatman® Nuclepore™ Track-Etched Membranes diam. 19 mm, pore size 0.05 μm). The mean diameter of the LUVs was verified by Dynamic Light Scattering (Zetasizer Nanoseries S, Malvern instruments). LUVs were always prepared freshly the previous day of the experiment.

To prepare the PDMS membranes, a TEM grid (G200H-Cu, Aname) was placed in the middle of the PDMS membrane ring. The membrane was subsequently plasma cleaned in a Harrick oxygen plasma cleaner using the following parameter: constant flow of oxygen between 0.4 to 0.6 mbar, high power, and exposure time between 15 and 60 s. A small 6 mm inner diameter ring was simultaneously plasma cleaned and bonded around the TEM grid. Then, the TEM grid was removed and the liposome solution was deposited and confined inside the thin bonded ring, with subsequent incubation for 1 h at room temperature. LUVs were then extensively washed with PBS buffer pH 7.5. The membrane was mounted in the stretching device placed in the microscope.

### FRAP of the pSLB

Patterned Supported Bilayers (pSLB) were obtained as described above on PDMS membranes, and the ring-containing membranes were mounted under an upright epifluorescence microscope (Nikon Ni, with Hamamatsu Orca Flash 4.0, v2). Images of pSLBs, obtained with either 15 s or 30 s plasma cleaning, were acquired with a 60x water dipping objective (NIR Apo 60X/WD 2.8, Nikon) and an Orca R2 camera. A small linear region of the pSLB was frapped by repeatedly scanning and focusing 180 fs pulses generated by a fiber laser (FemtoPower, Fianium) with central wavelength at 1064 nm at 20 MHz. A set of galvo mirrors (Thorlabs) and a telescope before the port of the microscope allowed to position and move (oscillations at 400 Hz) the diffraction limited spot at a desired place on the bilayer. Once bleached, fluorescence recovery was monitored for 5 min. Time-lapse imaging during the pSLB photobleaching and it recovery after photobleaching was done with a home-made software (Labview 2011).

### Sucrose loaded assay and Negative-stain Transmission Electron Microscopy

Sucrose loaded vesicles were prepared as previously described in the literature^17^, using a mixture of DOPC, DOPS and DOPE lipids in a 1:2:1 ratio. Lipids were evaporated and subsequently rehydrated with PBS buffer pH 7.5, containing 0.3 M sucrose. A solution of 0.6 mM lipids of vesicles was incubated for 20 min with 40 μM of Amphiphysin (non-fluorescent) at 37 C. The solution was incubated on a copper grid (G200H-Cu + Formvar, Aname), previously activated with 5 min UV) and subsequently stained with 2 % neutral phosphotungstic acid. Grids were imaged in a JEOL 1010 80kV TEM microscope.

### Mechanical/osmotic stimulation of the pSLB, protein injection and live imaging

Membrane-containing rings were mounted in the stretch system as previously described^7^. Image acquisition of cells and pSLBs were acquired with a 60x objective (NIR Apo 60X/WD 2.8, Nikon) in an inverted microscope (Nikon Eclipse Ti) with a spinning disk confocal unit (CSU-W1, Yokogawa), a Zyla sCMOS camera (Andor) and using the Micromanager software. The bilayer was stretched slowly for 120 s and the strain, obtained through the measurement of the hexagon extension, was between 5 and 8 %. After 120 s stretch, the bilayer was slowly released for 300 s. At release and upon tube appearance, images were acquired every sec in two different channels collecting each fluorophore emission signal. 3 μL of an amphiphysin or neutravidin stock solution (of a concentration depending on the desired end concentration but always in the same buffer as the one covering the pSLB to avoid any osmotic perturbation) was gently micro-injected in the buffer droplet hydrating the pSLB. End concentration ranged from 50 nM to 5 μM. In some instances, the non fluorescent protein was used to reach high concentrations. For the controls of tube behavior in absence of protein, no injection was performed. To modify osmolarity, the pSLB was exposed to medium mixed with de-ionized water and after pressurised cap formation, protein was injected in the same conditions as above. Osmolarity was adjusted to that of the buffer hydrating the pSLB.

### Supported Lipid Bilayer (SLB) formation on glass coverslips

SLBs on glass coverslips used for the calibration in the quantitative fluorescence microscopy were obtained as previously described^37^. Glass coverslips were cleaned by immersion in 5:1:1 solution of H2O:NH4:H2O2 at 65 °C for 20 min and were dried under a stream of N2 gas. GMVs were obtained as previously but with different lipid mixtures. To obtain SLBs with 0.1 to 0.5 % of protein-like fluorophores, 2 LUV-stock solutions were prepared, either DOPC only, or DOPC with 0.5 % 1,2-dioleoyl-sn-glycero-3-phosphoethanolamine-N-(TopFluor® AF488) (ammonium salt). Lipid films were rehydrated in 150 mM NaCl and 10 mM Tris, pH 7.4, to a final concentration of 3 mg/mL. GMVs were extruded as previously described to obtain LUVs. Small rings of 6 mm diameter of PDMS were bonded as described before using plasma cleaning of both substrates, forming a small chamber on top of the coverslip. Coverslips were activated by cleaning with oxygen plasma (Harrick) in a constant flow mode (pressure 0.6 and at high power XW) for 20 min. The two LUV stock solutions were diluted in fusion buffer (300 mM NaCl, 10 mM Tris, 10 mM MgCl2) to 0.5 mg/mL solutions at different ratios to obtain a set of solutions from 0 to 0.5 % TopFluor-AF488. SLBs of the different fluorophore ratios were obtained by incubating the diluted solutions in the glass coverslips chambers, immediately after the plasma cleaning process, for 1 h at room temperature. Liposomes were extensively rinsed with the fusion buffer and subsequently milli-Q water.

### Imaging of the SLBs, liposome and protein solutions on glass for quantitative fluorescence microscopy

SLBs on glass were imaged in the same condition as the pSLB on PDMS. For the AF-488 enriched SLB, the exposure time and laser power were the same as for the protein channel. For the LissRhod-DPPE enriched SLB, parameters were the same as for the lipid channel. Background for the AF-488 enriched SLB was obtained by focusing on a LissRhod-DPPE enriched bilayer and recording an image in the 488 nm channel. The opposite was done for LissRhod-DPPE enriched background. Fluorescence image of protein solutions at different concentrations, from 0 to 0.75 μM, and of LissRhod-DPPE enriched LUV solutions (from 0 % to 0.1 %) were recorded with the same settings as for the pSLB protein channel.

### Cell culture and transfection

Normal Human Dermal Fibroblasts derived from an adult donor (NHDF-Ad, Lonza, CC-2511) were cultured using Dulbecco’s Modified Eagle Medium (DMEM, Thermofisher Scientific, 41965-039) supplemented with 10 % FBS (Thermofisher Scientific, 10270-106), 1 % Insulin-Transferrin-Selenium (Thermofisher Scientific, 41400045) and 1 % penicillin-streptomycin (Thermofischer Scientific, 10378-016). Cell cultures were routinely checked for mycoplasma. CO2-independent media was prepared by using CO2-independent DMEM (Thermofischer Scientific, 18045-054) supplemented with 10% FBS, 1% penicillin-streptomycin, 1.5 % HEPES 1M, and 2 % L-Glutamine (Thermofischer Scientific, 25030-024). One day before experiments, cells were co-transfected with the membrane-targeting plasmid peGFP-mem and the pEGFP-C1-Amph1, Transfection was performed using the Neon transfection device according to the manufacturer’s instructions (Invitrogen). peGFP-mem was a kind gift from Pr. F. Tebar. and contained the N-terminal amino acids of GAP-43^38^, which has a signal for post-translational palmitoylation of cysteines 3 and 4 that targets fusion protein to cellular membrane, coupled to a monomeric eGFP fluorescent protein. pEGFP-C1-Amph1 was a kind gift of Pr. De Camilli and contained the full-length Amphiphysin 1 coupled to a mCherry fluorophore.

### Mechanical stimulation of the cells and live imaging

Cell mechanical stimulation was done as previously described^7^. Briefly, a 150 μL droplet of a 10 μg/mL fibronectin solution (Sigma) was deposited in the center of the membrane mounted in the ring. After overnight incubation at 4 C, the fibronectin solution was rinsed, cells were seeded on the fibronectin coated membranes and allowed to attach during 30 to 90 min. Then ring-containing membranes were mounted in the stretch system previously described^7^. Cell images were acquired with a 60x water dipping objective (NIR Apo 60X/WD 2.8, Nikon) and an Orca Flash 4.0 camera (Hamamatsu), in an upright epifluorescence microscope with the Metamorph software. Cells were always imaged in two different channels collecting each fluorophore emission signal, every 3 sec. They were imaged for 2 min at rest, 3 min in the 6 % stretched state (nominal stretch of the PDMS substrate) and 3 min during the release of the stretch.

### Quantifications

- **Diameter of tubes expelled by amphiphysin from sucrose loaded vesicles using TEM images** The diameter of the lipid tube reshaped by amphiphysin was measured using the TEM images from the sucrose loaded assay. Diameters at one or two places of tubes expelled from the vesicles were measured manually on 7 different high magnification images (*60k) of two independent experiments. The mean diameter was computed from these measurements.
- **Binding curves of the protein to the buds and tubes** Stacks of the acquired images were prepared in Fiji. A stack containing a single lipid object (tube or bud) was isolated from the timelapse stacks obtained in protein channel, as well as a stack of a small area of the pSLB close to the object. Objects were automatically thresholded in CellProfiler and their mean florescence intensity was extracted. After background correction, the fluorescence intensity was plotted over time for each object.
- **Protein enrichment on the reshaped tube** The raw intensities of the elongated tubes were measured as explained above (in the tube diameter section), for both lipid and protein channels, at the same timepoint. The raw intensity of the bud in both channels was also measured assuming a spherical shape. We define the tube versus bud enrichment in both channels by the ratio between the mean intensities of the tube and bud. Mean intensities are calculated by dividing the raw intensities by the area of the tube or bud, which is the same in both channels. In the case of the lipid image, no enrichment is assumed. We thus normalize the protein enrichment value with that of the lipid which makes our measurement independent of geometry. See also Suppl Fig. 5a.
- **Estimation of protein coverage** To estimate the coverage of tubes and buds with amphiphysin, we first prepared flat membrane bilayers containing 0.5 % Liss Rhodamine fluorofore, and measured their average fluorescence intensity per unit area. Then, tubes or buds in experiments were identified as described in the “binding curve method”, and their average fluorescence intensity in the lipid channel was also calculated. By calculating a ratio between both values, we obtain a geometrical correction factor. Due to the 3D shapes of tubes and buds, this accounts for loss of signal if not all fluorescence is collected in the confocal slice, or gain of signal due to integration of fluorescence due to the 3D object. Then, we prepared flat membrane bilayers, but labelled with the same fluorophore used for amphiphysin, AF-488. By measuring florescence intensities as a function of AF-488 concentration, we obtained a calibration curve between the fluorescence signal and fluorophore concentration, as previously described^37,27^. Finally, we measured the average fluorescence intensity of tubes and buds in the amphiphysin channel, and used the calibration curve and the geometrical correction factor to estimate an amphiphysin dimer concentration (accounting for the number of fluorophores per dimer). After assuming a dimer area of 58 nm2 (same area as in our simulations, close to the one classically used^4^), we finally obtain a coverage estimation (see also Suppl. Fig. 7a).
- **Quantification of amphiphysin tubulation in the cell experiments** In videos of Amphiphysin over-expressing cells, time slots of 90 s before, during and after stretch were analysed. The number of tubulations appearing in each one of the slots was manually counted having as reference the timepoint of formation of the structure.
- **Quantification of the elongation rate** For both tubes elongating from buds in-vitro or tube elongating in the cellular plasma membrane, the elongation rate was obtained by plotting the length of the tube (increasing with time) at different time points. The slope of the fit to a linear curve directly gives the elongation rate.

## Theoretical Model

### Introduction

In previous work ^23^, we presented a general continuum framework to study the dynamics of curved protein-membrane interaction, highlighting the role of protein curvature and self-interaction on their curvature-sensing and generation capability. This theory accounted for protein diffusion and for membrane elasticity and hydrodynamics. Here, we extend it to include the sorption dynamics of proteins ^39^ and their orientational order ^21^. Indeed, some curved proteins such as those containing BAR domains are elongated objects interacting anisotropically with the curvature of the membrane. With this extended theory, we can predict the coupled dynamics of protein area fraction *ϕ*, the shape of the membrane surface Γ parametrized by ***x***(*u, v, t*), and nematic order. Orientational order is quantified by the traceless and symmetric tensor

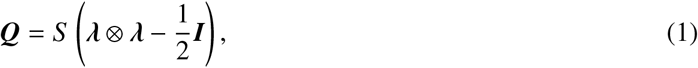

capturing two important pieces of information: the order parameter *S* taking values between 0 and 1, where 0 corresponds to an isotropic organization of proteins and 1 to the maximum degree of order, and the net protein orientation given by the unit vector ***λ. I*** is the identity tensor on the surface. In the following, we denote by ***k*** the second fundamental form or curvature of the surface, characterizing the curvature of the surface along any given direction.

### 1 Modeling the state prior to protein exposure

Prior to protein exposure, we model the formation of membrane protrusions following the conceptual and computational approach in ^6^ under the assumption of axisymmetry. We consider an inextensible membrane patch Γ of radius 1 μm interacting with a support through an interaction energy density per unit surface area *U*(*z*), where *z* is the separation between the membrane and the substrate. The free energy of the membrane is

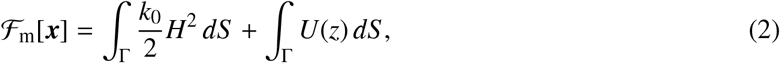

where *k*_0_ is the bending stiffness of the membrane and *H* = (tr ***k***)/2 is the mean curvature and *dS* is the area element of Γ. To model the dynamics and as described elsewhere ^23,40,41^, we introduce a dissipation potential accounting for membrane viscosity *D_m_*[***v***], and obtain the governing equations of the system by minimizing the Rayleighian functional

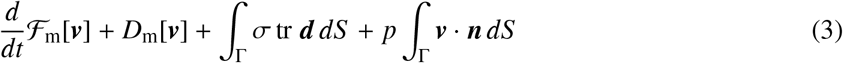

with respect to the membrane velocity **v**, where ***d*** is the rate-of-deformation tensor of the membrane, ***n*** is the outer normal to the surface, and the surface tension *σ* and pressure *p* are Lagrange multipliers that enforce the local inextensibility of the membrane and global incompressibility of the fluid enclosed between the membrane and the substrate.

Excess membrane resulting from lateral compression and excess enclosed volume resulting from osmotic imbalances lead to a variety of equilibrium membrane protrusions, which include long tubules, spherical buds and shallow spherical caps as mapped in ^6^. All of these protrusions are observed in our experimental system. To study the effect of proteins on each of these types of structures, we prepared protrusions in equilibrium by laterally compressing the flat membrane and increasing the enclosed volume *V* with respect to *V*_0_, the reference volume for a planar membrane at the equilibration separation *z*_0_. Upon compression, the membrane delaminates to form a shallow spherical cap. By further increasing lateral compression and/or the enclosed volume, we obtained equilibrium structures consisting of spherical buds connected to the supported part of the bilayer by a narrow neck, or long tubular protrusions, see Fig. 2g in main text.

### 2 Modeling the protein-membrane interaction dynamics

Proteins on the membrane are represented by two fields, their area fraction *ϕ* and their nematic tensor ***Q***, which in our axisymmetric setting can be represented by the order parameter *S* and the angle *θ* with respect to the azimuthal direction.

Given the free-energy of the elongated and curved proteins on the membrane 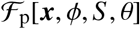, to be specified later and which depends on ***x*** through the curvature of the membrane, we can define the total free energy of the system as

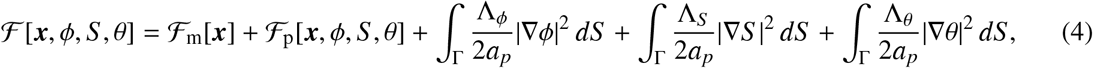

where the last three terms regularize the phase boundaries between regions of different protein coverage, order and orientation.

We can write the Rayleighian functional of the membrane-protein system as

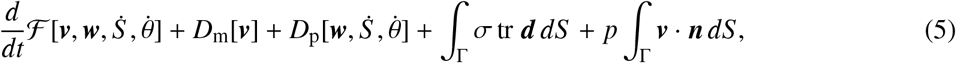

where ***w*** is the net diffusive velocity of proteins relative to the lipids and 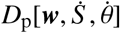 is the dissipation potential of proteins accounting for drag due to changes in position and nematic order of the proteins. Minimization of the Rayleighian with respect to ***v*** leads to the equations of mechanical equilibrium govering shape dynamics and lipid flow. Minimization with respect to ***w*** leads to a generalized Fick’s law relating ***w*** to the gradient of the chemical potential of the proteins, whereas minimization with respect to 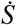 and 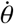 leads to configurational balance equations. Here, we assume that S and θ relax much faster than ***x*** and *ϕ*. Combining Fick’s law with the equation of balance of mass for proteins

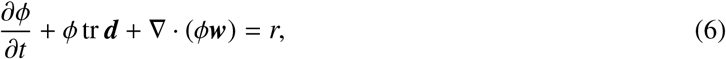

where *r* is the sorption rate, we obtain a nonlinear diffusion-reaction equation for the protein density. All these equations are self-consistently coupled in this formalism. We refer to ^23^ for a full account of this formulation and of its computational axisymmetric implementation using a Galerkin finite element method based on B-Spline approximations.

We model sorption with a modified Langmuir model given by ^39^

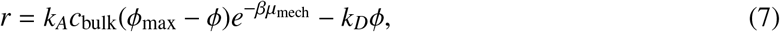

where *k_A_* is an adsorption rate constant, *c*_bulk_ the bulk concentration of proteins, *μ*_mech_ is the mechanical part (associated with their bending elasticity) of the chemical potential of proteins on the membrane, explicitly defined in Eq. (10), 1/β is the thermal energy, and *k_D_* is a desoprtion rate constant. The exponential part of the adsorption term models an adsorption mechanism by which a curved molecule in solution must conform to the membrane curvature by a thermal fluctuation to become a membrane-bound protein, analogously to the case of binding of flexible adhesion molecules ^42^. This has kinetic and thermodynamic consequences, as adsorption becomes faster and equilibrium coverage higher when the membrane curvature is close to the spontaneous curvature of the protein.

### 3 Free-energy of the elongated and curved proteins on a membrane

To our knowledge, previous theoretical continuum models for the free-energy of curved and elongated molecules on membranes cannot predict the simultaneous evolution of membrane shape and nematic order, and instead fix nematic order and direction ^43,44^. To understand the interaction between elongated curved proteins with a membrane and interpret our observations, we developed in a companion paper ^21^ a new mean field density functional theory accounting for protein area coverage, orientational order and membrane curvature, which we summarize next.

#### 3.1 Mean field density functional theory

The theory used here is an adaptation and generalization of that recently proposed by ^22^ for hard ellipsoidal particles, which corrects Onsager’s classical theory of isotropic-to-nematic transitions for non-spherical particles to provide quantitative prediction at high densities and moderate particle aspect ratio. Our extension of the theory also accounts for the curvature energy of the proteins adsorbed on a curved surface. Following a mean field approximation and a passage to the continuum limit, the free-energy of the ensemble of elongated molecules is expressed in terms of the position-dependent number density of proteins *ψ*, related to the area fraction by *ϕ* = *a_p_ψ* where *a_p_* is the area of a protein, and the angular distribution *f* of proteins as

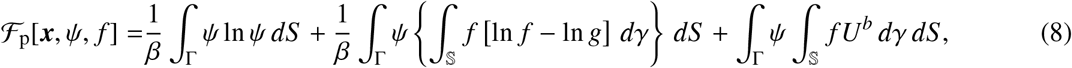

where 1/*β* = *k_B_T* is the thermal energy and the set 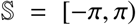 represents all possible orientations of molecules. The first term models the positional entropy of proteins, the second term accounts for orientational entropy and the excluded area though the function *g*(*ψ, γ*) = 1 - *ψ*[*c* – *dS P*_2_(cos *γ*)] (with *c* and *d* parameters that depend on the geometry of the particles and *P*_2_(*x*) = *x*^2^ - 1/2), and the last term models the bending elasticity of proteins. The function *U^b^*(***k***, *γ*) is the bending energy of an adsorbed protein oriented along the tangential vector ***i*** forming an angle γ with a fixed direction and is given by

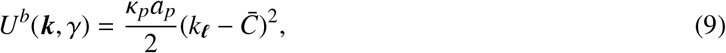

where *κ_p_* is its bending rigidity (with units of energy), 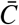 is its preferred curvature along the long axis, and *k_i_* = ***ℓ · k · ℓ*** is the normal curvature of the surface along the long direction of the protein. Equation (8) allows us to identify the mechanical part of the chemical potential of proteins as

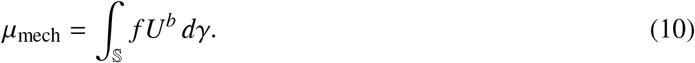

Minimization of 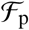 with respect to the angular distribution *f* yields an effective free energy depending only on the nematic tensor

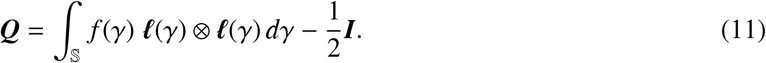

Since ***Q*** is traceless and symmetric, it can be expressed as in Eq. (1). In the axisymmetric setting considered here, ***Q*** can be parametrized by *S* and the angle *θ* between the nematic direction ***λ*** and the azimuthal direction. Denoting by *k*_1_ and *k*_2_ the principal curvatures of the surface at any point (which in the axisymmetric setting considered here are along symmetry directions,), we can express the free energy of the proteins as

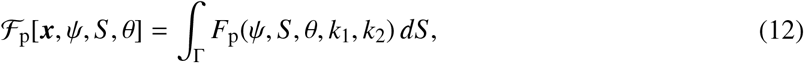

where the evaluation of *F*_p_(*ψ, S, θ, k*_1_, *k*_2_) involves the solution of a nonlinear system of algebraic equations with two unknowns, see ^21^. Importantly, the only material parameters in this theory are the long and short axes of the ellipse modeling a protein, its preferred curvature 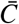 and its bending stiffness *κ_p_a_p_*, for which estimates are available.

This theory allows us to evaluate the free energy of proteins and study the isotropic-to-nematic transition on membranes adopting simple geometric motifs observed in our experiments, such as spheres and cylinders of various radii. On spheres, the free energy above is independent of *θ* whereas on cylinders it is minimized for *θ* = 0 (proteins aligned with the direction of curvature) as long as the cylinder radius is larger than 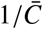. Minimization with respect to *θ* allows us to compute the free-energy profile as a function of area coverage *ϕ* and order *S* alone, see Fig. 2 and Supp. Fig. 3. These figures shows that the free-energy landscape exhibits an order- and coverage-dependent forbidden region due to crowding effects. It also shows that, for a planar and a spherical configuration, the model predicts a sharp and discontinuous isotropic-nematic phase transition with a range of intermediate protein coverages exhibiting coexistence of the two phases. The landscape on cylindrical surfaces is different in several ways. There, the isotropic-to-nematic transition is continuous and the isotropic phase is ordered even at low *ϕ*, particularly for thin tubes, due to the bias introduced by anisotropic curvature. See ^21^ for further details.

#### 3.2 Explicit parametrization of the theory

This mean field theory connects the microscopic statistical physics with continuum physics and predicts the density- and curvature-dependent isotropic-to-nematic transition of proteins, but is cumbersome to evaluate and integrate in the computational framework described in Section 2 and in ^23^. For this reason, we fit the free energy of proteins given by the mean field theory, 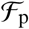, to an explicit functional form that we denote as 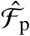. Replacing 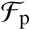 by 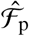 is done for purely practical reasons. To identify an ansatz for the functional form of 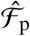, we examine Eq. (8).

Focusing first on the first two integrals in this equation, which do not depend explicitly on *θ*, and noting that 〈*P*_2_(cos *γ*)〉 = *S* /2 where 〈 〉 denotes the average with respect to *f* ^21^, we postulate the entropic part of the ansantz as

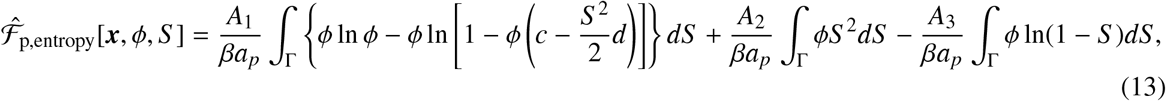

where *A*_1_, *A*_2_ and *A*_3_ are non-dimensional fitting coefficients. The first integral accounts for positional entropy and excluded area, the second integral is a quadratic approximation to the order entropy, and the last integral allows us to fit the fast increase in the free-energy landscape for large *ϕ* and *S*, Supp. Fig. 3a1,b1.

Focusing now on the last term of Eq. (8), to propose an explicit functional form for the curvature energy of proteins, we note that

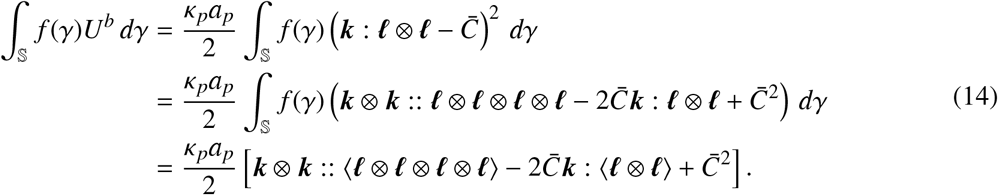

where : denotes the double contraction of second-order tensors, :: the contraction of fourth-order tensors, and 〈 〉 the average with respect to *f*. We note that 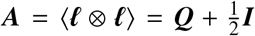. Invoking the Doi closure ^45^ according to which 〈***ℓ*** ⊗ ***ℓ*** ⊗ ***ℓ*** ⊗ ***ℓ***) :: ***C*** ≈ 〈***ℓ*** ⊗ ***ℓ***) ⊗ 〈***ℓ*** ⊗ ***ℓ***) :: ***C*** where ***C*** is a fourth-order tensor, we can approximate the curvature part of the proteins free energy as

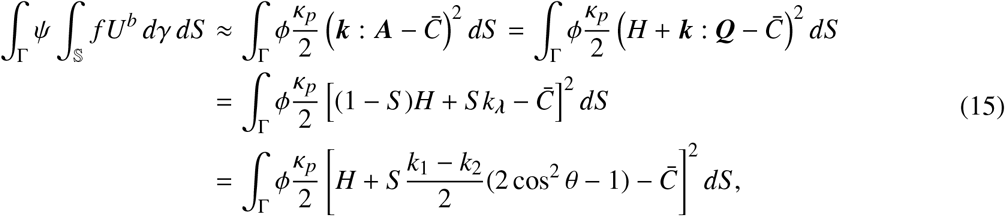

where *k_λ_* = ***λ*** · ***k*** · ***λ*** is the normal curvature along the nematic direction. We checked that this approximation to the curvature part of the free energy was insufficient to closely fit the free energy of cylinders (particularly the minimum energy paths and isotropic-to-nematic transition in Supp. Fig. 3a) and for this reason we consider an expanded ansantz for the curvature free energy of proteins of the form

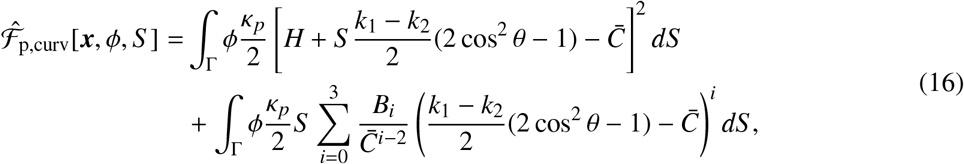

where *B_i_* are nondimensional fitting parameters. We note that the new term in the second line only adds a constant unless curvature is anisotropic. Combining Eqs. (13,16), we obtain an explicit form of the protein free-energy functional 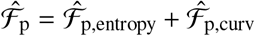 approximating the mean field functional 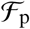 and amenable to numerical calculations.

To fit the parameters *A_i_*, we first focused on the purely entropic interaction of elliptical proteins on a flat membrane, which we evaluated with the mean field model in Eq. 8. We then fitted the functional proposed in Eq. 13 to the mean field landscape using a nonlinear least-squares method. In a second step, we included the bending energy in the mean field model, computed the free-energy landscape for flat, spherical and cylindrical configurations with different curvatures, see Supp. Fig. 3a, and used these free-energy landscapes to fit *B_i_* using nonlinear leasts-squares.

See Supp. Fig. 3 for a comparison of the free-energy profiles obtained with both models. Although there are noticeable differences, these are small and the approximate functional 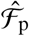 captures quantitatively the most salient features of the mean field model including the curvature-dependent isotropic-to-nematic transition.

We note that if the radius of curvature of a cylindrical membrane is larger than that of the protein 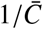, then the free energy is minimized for *θ* = 0 ^21^. Thus, although we fitted the mean fit model to include the smaller radii, in all our simulations radii of curvature were larger than 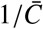, and thus the free-energy functional can be simplified by setting 2 cos^2^ *θ* – 1 = 1.

### 4 Selection of parameters

Following ^6^, we assume a Morse potential for *U*(*z*) with a membrane-support equilibrium distance of *z*_0_ = 4.4 nm and adhesion energy −*U*(*z*_0_) ≈ 1.5 mJ/m^2^. For the material properties of the lipid membrane, we consider *k*_0_ = 20 *k_B_T* for the bending stiffness and *η* = 5 · 10^-9^ Nsm^-1^ for the 2D viscosity.

We consider N-BAR proteins to be elliptical with semi-axis lengths *a* = 7.5 nm and *b* = 2.5 nm, leading to the non-dimensional constants *c* = 15.66 and *d* = 6 appearing in the expression for the free-space as a function of density and order in Eq. (13). We assume that these proteins have an intrinsic curvature of 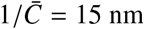, an area on the membrane of *a_p_* ≈ 58 nm^2^, and a protein bending rigidity of *ϕ_max_k_p_* = 20 *k_B_T* at saturation (*ϕ_max_* ≈ 0.75) based on a rigidity of the membrane-protein compound of 40 *k_B_T* ^18,20,29^. We consider a diffusion coefficient for proteins of *D*_p_ = 0.13 μm^2^/s.

The fitting procedure of the functional given by Eqs. (13,16) results in *A*_1_ = 1.25, *A*_2_ = 0.7, *A*_3_ = 0.5, *B*_0_ = 1.61, *B*_1_ = −2.49, *B*_2_ = 1.32 and *B*_3_ = −0.43. We consider an adsorption rate of *k_A_* = 1/30 μM^-1^ s^-1^ and desorption rate to *k_D_* = 1/1800 s^-1^, in the order of that consider in previous works in a related system ^4^.

In the absence of measurements, we choose Λ_*ϕ*_/*a_p_* = 10k*_B_T* and Λ_*S*_ /*α_p_* = Λ_*θ*_/*a_p_* = 1*k_B_T* large enough so that, when phase separation occurs, domain boundaries have a finite thickness and simulations are devoid of numerical oscillations signal of ill-conditioning, and small enough so that the dynamics of the problem are not affect by these parameters.

Our chemo-mechanical model for protein-membrane interaction captures many different phenomena occurring at different time-scales. To interpret our simulations and understand the observed dynamics, we examine next the timescales of the major phenomena. We have already mentioned that we assume that orientational order relaxes very fast. We approximate the time required for membrane shape dynamics as 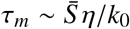 where 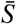 is the typical surface area of a geometric feature ^46^, leading to the estimate *τ_m_* ≈ 0.01 s. The timescale for protein diffusion is *τ_p_* ~ *ℓ*^2^*/D_p_* where *ℓ* is either the radius of a spherical bud or the length of a tube where proteins diffuse. For the membrane protrusions studied here, we estimate *τ_p_* to be a few seconds. The timescale for protein adsorption can be approximated as *τ_A_* ≈ 1 /*k*_*A*_*c*_bulk_, which in our experimental conditions ranges between a few seconds to minutes depending on concentration. Thus, even if these estimates are crude, the system exhibits a significant scale separation, except at high protein concentrations where protein adsorption and diffusion may compete. As discussed in the main text, in experiments there is another time-scale associated to bulk transport of proteins in the medium, not accounted for in the model.

### 5 Simulation protocol for protein-membrane interaction

To computationally examine the effect of BAR proteins on pre-existing membrane protrusions, we start from tubular or spherical protrusions in mechanical equilibrium as those in Fig. 2g and following the protocol described in Section 2 of this supplement. Then, we prescribe the protein concentration in the medium, *c*_bulk_, and computationally track the dynamics of the system as described in Section 3 of this supplement with the explicit free energy of proteins described in Section 4.2 of this supplement. According to Eq. (7), protein adsorption is faster in highly curved regions of the membrane, such as necks of spherical buds. Furthermore, curvature gradients also generate gradients of chemical potential that drive protein diffusion on the membrane towards highly curved regions. As these regions, with possibly anisotropic curvature, become enriched, nematic order progressively develops, giving rise to the protein dynamics and reshaping described in the main text.

In the actual system with many membrane protrusions interacting with proteins, the reshaping of one protrusion may result in lipid and enclosed water exchange with the rest of the system, in particular with the adhered part of the membrane surrounding it. Since in our computational model we study one protrusion in isolation, we need to specify a mechanical ensemble controlling lipid and enclosed volume exchange between a protrusion and the adhered membrane. In our simulations, we consider an inextensible and axisymmetric membrane patch. During dynamical simulations, we fix the volume enclosed between the membrane patch and the substrate to its initial value. At the boundary of the patch, we impose a constant membrane tension given by the membrane tension in the equilibrium state prior to protein exposure. Thus the edge of the patch can move to accommodate lipid exchange between the protrusion and the adhered part of the membrane. The protrusion can also exchange water with the adhered part of the system, either because the later is changing its size or because the membrane is changing its separation *z* with the substrate. We found that modifying the mechanical ensemble, e.g. fixing the projected area of the patch instead of tension or pressure difference instead of enclosed volume, had some effect on the dynamics but did not fundamentally modify the protein dynamics and reshaping mechanisms described here. With our ensemble, however, elongation of the neck of a bud did not lead to significant shrinkage of the spherical bud, as observed in many experimental instances, but rather to membrane exchange between the protrusion and the adhered part (Supp. Video 7). We identified that this was due to the limited ability of the protrusion to expel enclosed volume underneath the adhered part in our simulations, and when this volume exchange was eased, for instance by considering a more compliant membrane-substrate interaction, we recovered the experimental phenomenology of bud consumption upon neck elongation (Supp. Video 8).

## Acknowledgements

This work was supported by the Spanish Ministry of Science and Innovation (PGC2018-099645-B-I00 to X.T., BFU2016-79916-P and PID2019-110298GB-I00 to P. R.-C.), the Spanish Ministry of Economy and Competitiveness/FEDER (BES-2016-078220 to C.T., the European Commission (H2020-FETPROACT-01-2016-731957), the European Research Council (CoG-616480 to X.T., CoG-681434 to M.A.), the Generalitat de Catalunya (2017-SGR-1602 to X.T. and P.R.-C., 2017-SGR-1278 to M.A), the prize “ICREA Academia” for excellence in research to P.R.-C. and to M.A., Fundació la Marató de TV3, and Obra Social “La Caixa”. IBEC and CIMNE are recipients of a Severo Ochoa Award of Excellence from the MINECO.

We would like to thank all the members of P. Roca-Cusachs, X. Trepat and M. Arroyo laboratories for technical assistance and discussions. We thank M. Pons, X. Menino, M.G. Parajo, M-A. Rodriguez, N. Castro, R. Sunyer, the Unitat de Criomicroscòpia Electrònica (Centres Científics i Tecnològics de la Universitat de Barcelona, CCiTUB), and the MicroFabSpace and Microscopy Characterization Facility, Unit 7 of ICTS “NANBIOSIS” from CIBER-BBN at IBEC, for their excellent technical assistance.

## Contributions

A.L.LR, C.T., N. W., M.A and P.R.-C. conceived the study; AL.LR, X.Q., M.S., X.T. and P.R-C designed the experiments, C.T, N-.W. and M.A designed the simulation; AL.LR., X.Q., D.Z and M.S. performed the experiments; C.T and N-.W. performed the simulation. AL.LR, X.Q., and P.R-C. analyzed the experiments; AL.LR, C.T, N-.W. and M.A analyzed the simulation; and A.L.LR., M.A. and P.R-C. wrote the manuscript. All authors commented on the manuscript and contributed to it.

## Competing interests

The authors declare no competing financial interests.

---

**Suppl. Figure 1:**
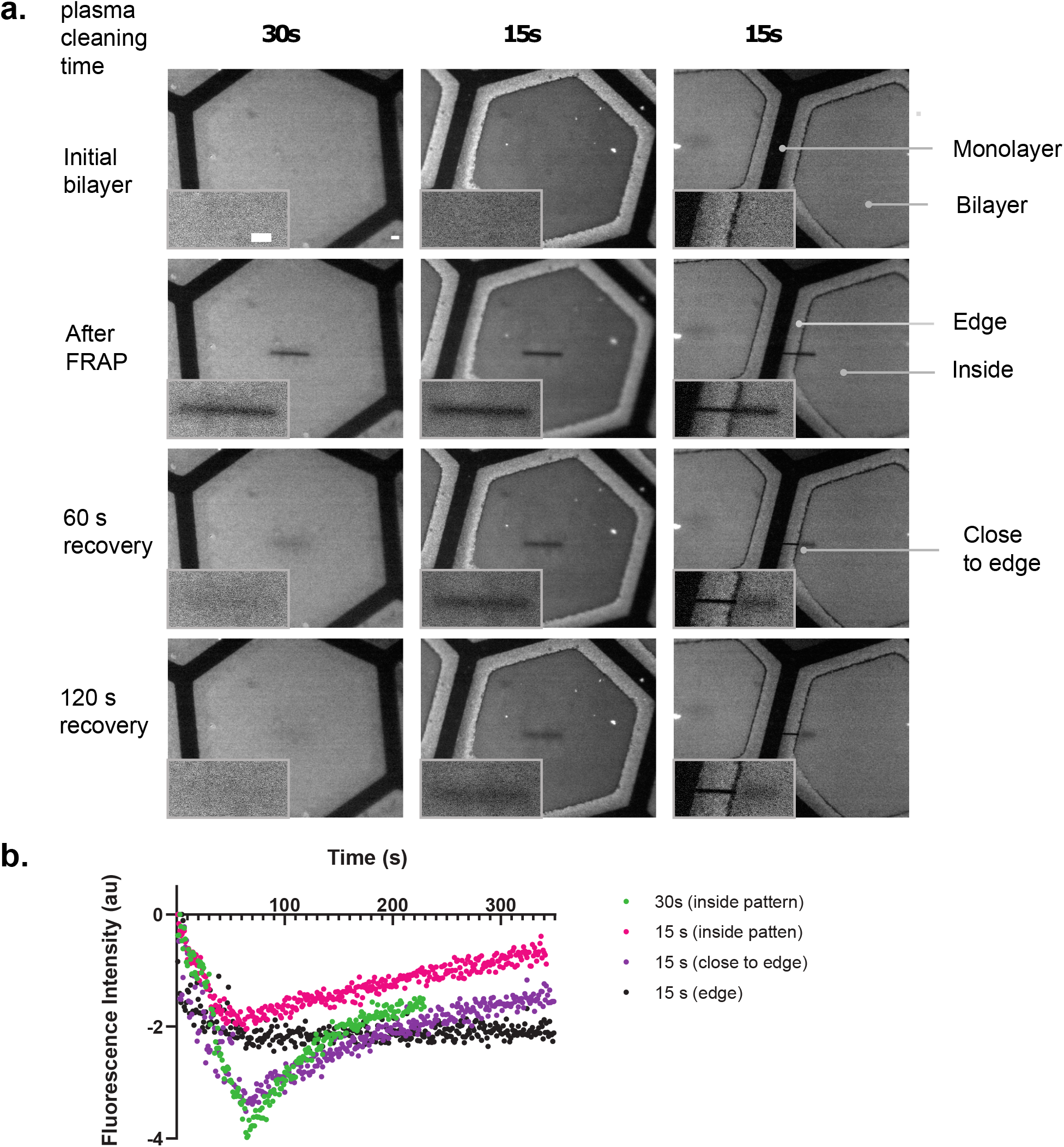
Characterization of membrane fluidity. **a**, Time-lapse images of the Patterned Supported Lipid Bilayers (pSLB) obtained after photobleaching a line. Results after different plasma cleaning times of the PDMS membrane are shown. Shorter plasma cleaning time (15 s) lead to more liposomes sitting on top of the bilayer and at the edge of the hexagon. Fluorescent recovery after photobleaching (FRAP) experiments show that both pSLBs (15 s or 30 s plasma cleaning times) are fluid compared with the non-fluid border (right images). Scale bar, 5 *μ*m. **b**, Recovery curves of the frapped areas of several pSLBs obtained either with a 15 s or a 30 s plasma cleaning time. Recovery is slower with a shorter plasma cleaning time, indicating a lower membrane fluidity. The edge does not show recovery.

**Suppl. Figure 2:**
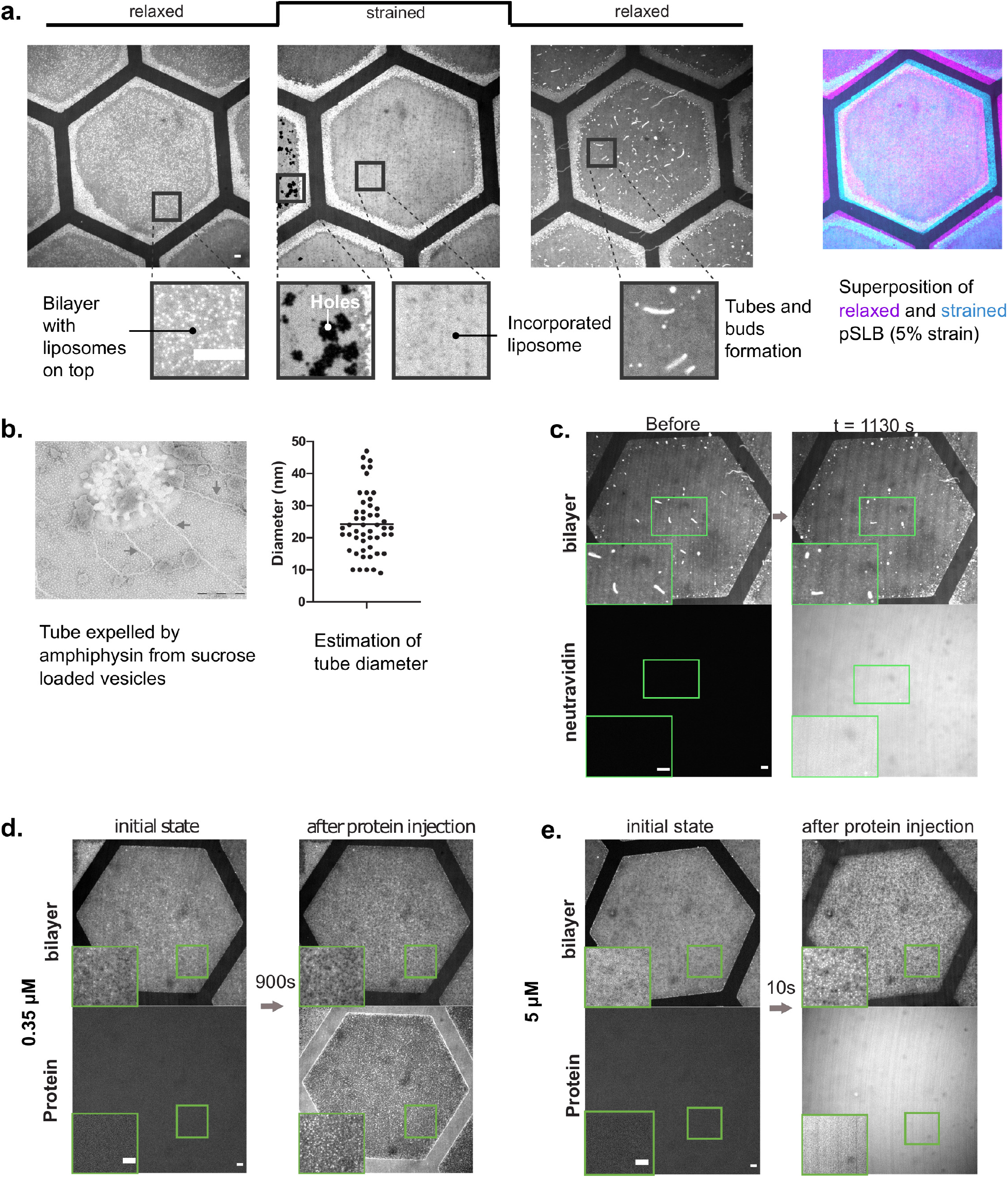
Additional characterization of membrane reshaping. **a**, (Left) Detailed example of the formation process of tubes and buds. At the relaxed initial state, liposomes stand in excess on top of the Patterned Supported lipid Bilayer (pSLB). With strain, the liposomes incorporate in the pSLB, and if not enough excess lipid is present, holes are formed and the naked PDMS membrane is exposed (dark holes). Upon release, excess lipids are expelled in the form or tubes or buds. (Right) Superposition of the relaxed bilayer (magenta) and strained bilayer (cyan). **b**, Estimation of the diameter of tubes reshaped by Amphiphysin using vesicles incubated with the protein and subsequently observed by transmission electron microscopy (TEM). **c**, Control in which 1 *μ*M fluorescent neutravidin is injected on top of the pSLB. The buds remain intact and the tubes slowly relax to buds, but no reshaping in the form of thin tubes is observed. **d, e**, Example of Amphiphysin injected on top of a non-stimulated pSLB at low (d) and very high (e) concentrations. At low concentration, no major effect is observed. At very high concentration, the bilayer is teared by the protein, leading to an immediate pSLB reshaping, in the form of bright dots and black holes. Scale bars, 5 *μ*m.

**Suppl. Figure 3:**
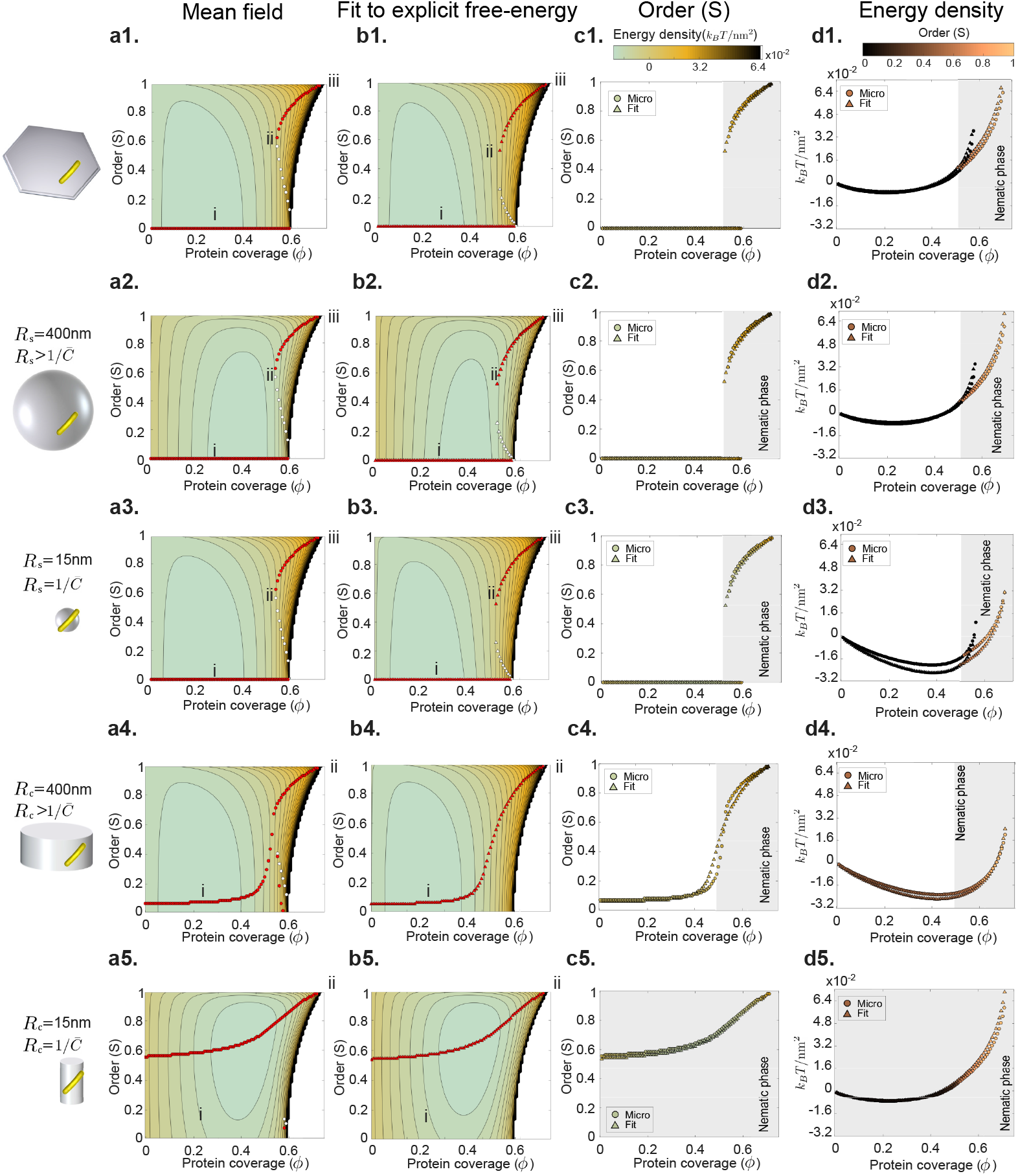
Explicit parametrization of the mean field model. **a**, Landscape of the free-energy density computed with the mean field model in Eq. (11) in the Theoretical Model and described in detail in ^21^ for membranes of different curvature (flat, spherical and cylindrical with radii larger or equal to the intrinsic radius of a protein 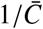). **b**, Analogous landscapes of the free-energy density with the explicit model given by Eqs. (12,15) fitted to the mean field model. This explicit model 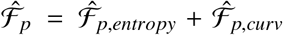 approximation is amenable to numerical calculations, see Theoretical Model. By minimizing the free-energy density with respect to S for a given protein coverage *ϕ* we find equilibrium paths *ϕ*(S). Stable branches are marked with red dots and unstable ones by white dots in (a) and (b). **c**, Comparison of the stable branches in the *ϕ* - S plane with both models (color is energy density). **d**, Comparison of the stable branches in the *ϕ* - energy plane with both models (color is order).

**Suppl. Figure 4:**
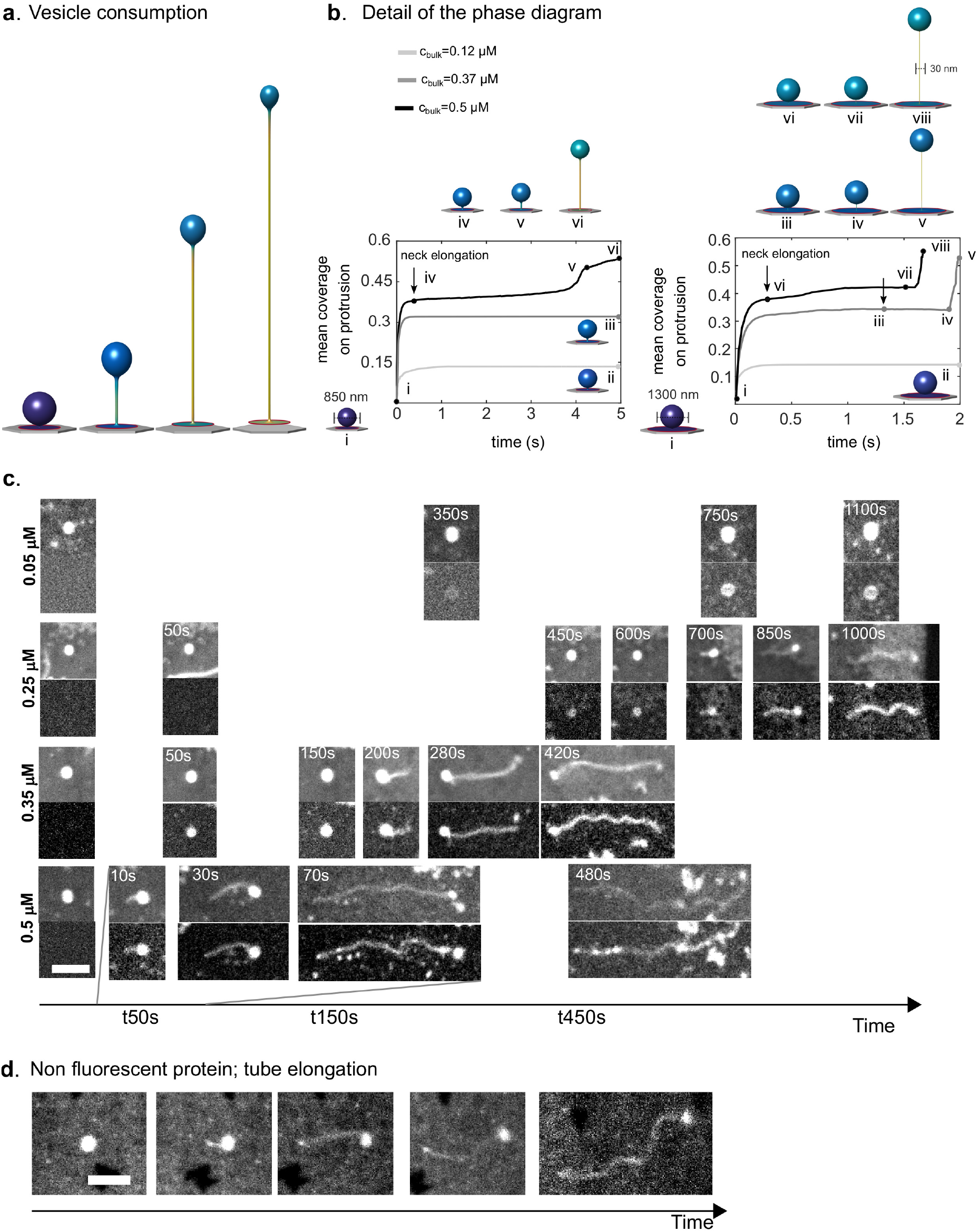
Time-and concentration-dependent bud reshaping. **a**, Simulation of a bud of 850 nm diameter exposed to a concentration of 0.5 *μ*M where exchange of the volume enclosed by the protrusion is eased by considering a softer substrate interaction, see Theoretical Model. Tube elongation is concomitant with consumption of membrane area of the vesicle. **b**, Examples of the numerical simulations used to build the dynamical diagram where two buds of different initial radius are exposed to a set of protein bulk concentrations. Membrane tension is fixed to that prior to protein exposure and the volume enclosed between the membrane (protrusion and adhered part) and the substrate is fixed. Membrane reshaping is faster at higher concentration and a threshold protein coverage is required for neck elongation. **c**, Experimental examples of buds elongated from their neck, at different concentrations of Amphiphysin in the bulk. Buds are elongated faster at higher concentration. d, Example of a bud elongated by non-fluorescent Amphiphysin. Scale bars, 5 *μ*m.

**Suppl. Figure 5:**
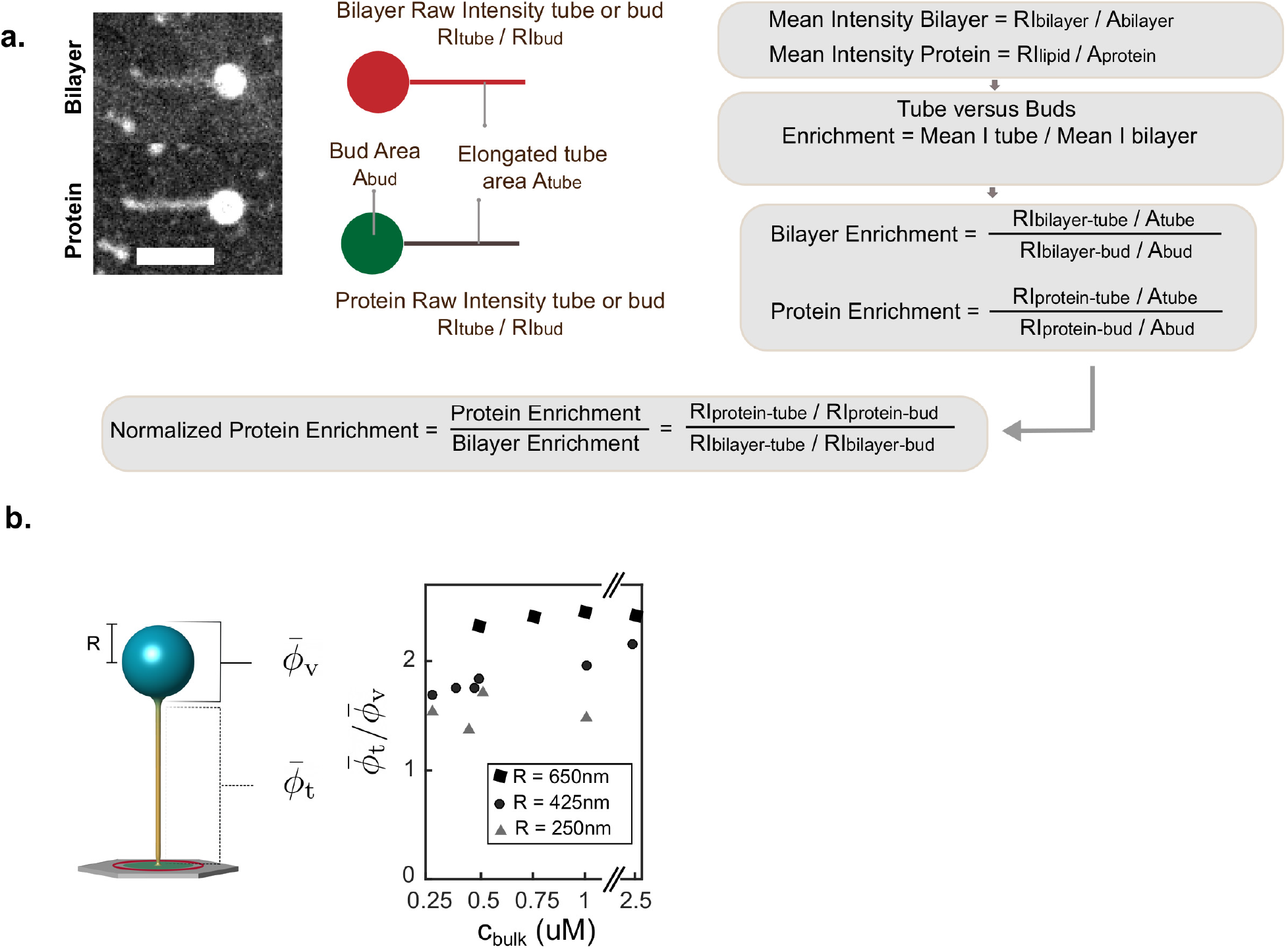
Estimation of Amphiphysin enrichment. **a**, Quantification of protein enrichment on the elongating tube. (Left) Examples of bilayer and protein fluorescence images of an elongating tube. Raw intensities on the tube and bud are measured (and corrected from background) in both lipid and protein images at the same timepoint. (Right) Protein enrichment on the tube versus bud is defined as the ratio between the mean protein fluorescence intensity levels in tubes and buds. However, calculating mean protein intensities require calculating membrane areas, which is challenging. To circumvent this, we assume that real enrichment in the membrane bilayer channel is 1, that is, the concentration of membrane is the same in both tubes and buds. Then, we factor out membrane areas by normalizing enrichment in the protein channel by the same value in the membrane bilayer channel. **b**, Computational estimation of relative protein enrichment between mean coverage on the tube 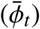 and mean coverage on the vesicle 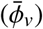 for buds of different sizes exposed to different protein concentrations.

**Suppl. Figure 6:**
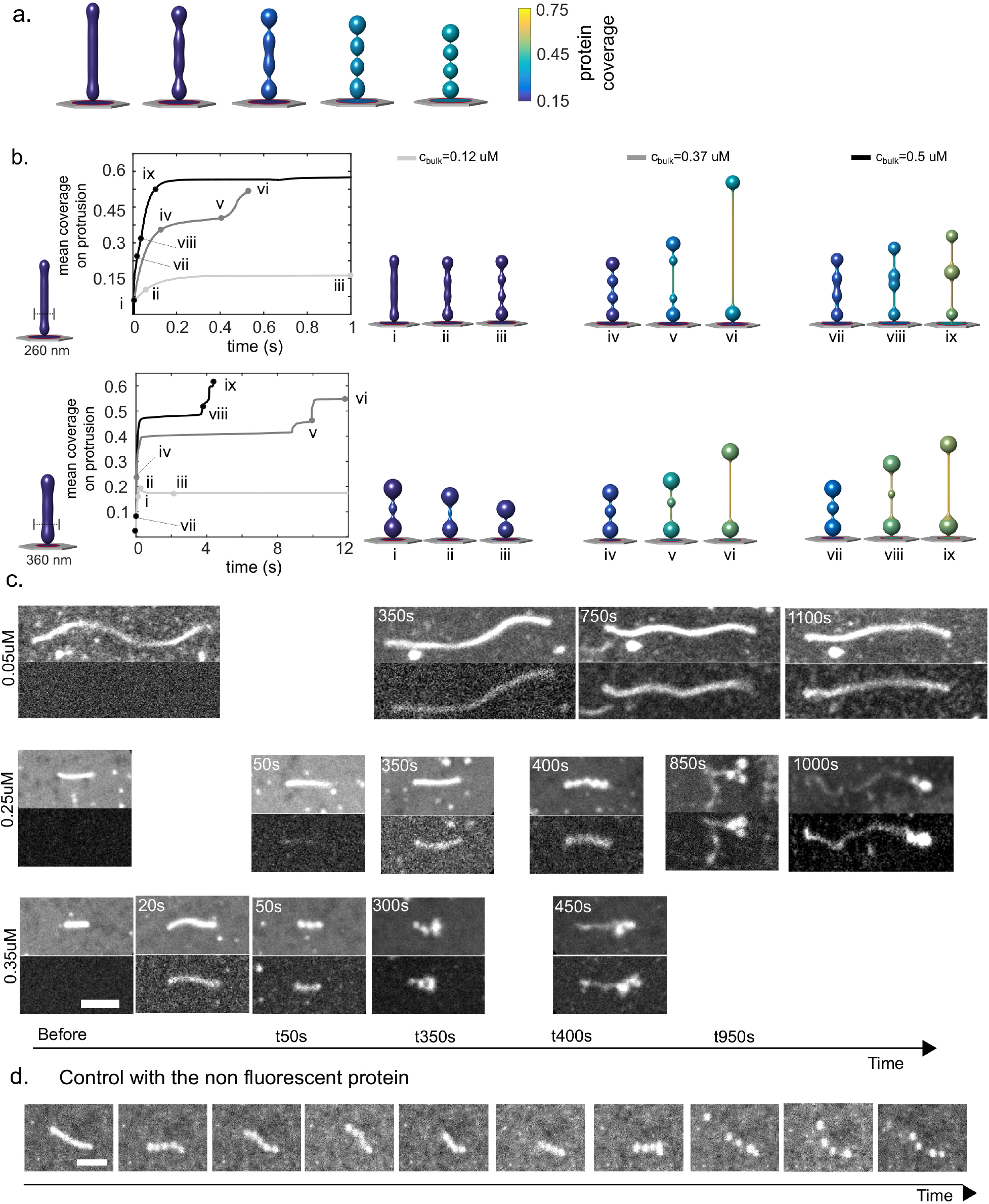
Time-and concentration-dependent tube reshaping. **a**, Membrane reshaping in the absence of a nematic transition (by prescribing isotropic orientational order, S=0); as predicted in the mean field theory, the saturation protein coverage is ≈ 0.55. **b**, Examples of the numerical simulations used to build the dynamical diagram where two tubes of different initial radius are exposed to a set of protein bulk concentrations. Membrane reshaping is faster at higher concentration. While proteins bind, the tubes connecting the pearl are shrinking. **c**, Experimental examples of tubes reshaped at different concentrations of Amphiphysin in the bulk. Pearling and elongation occur faster at higher concentration. **d**, Example of a tube undergoing the pearling phase due to binding of non-fluorescent Amphiphysin. Scale bars, 5 *μ*m.

**Suppl. Figure 7:**
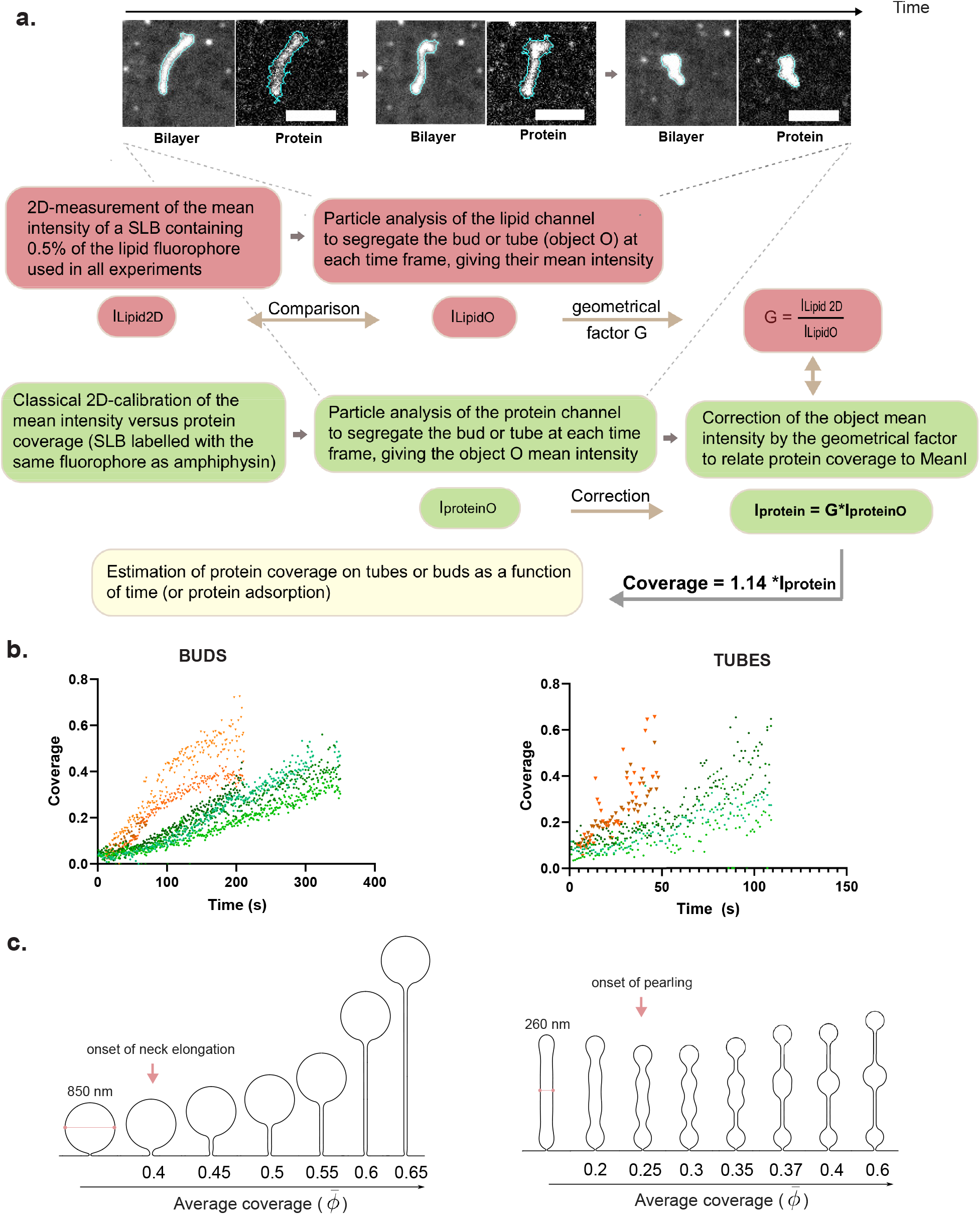
Estimations of protein coverage. **a**, (Top) Representative images of the particle analysis performed at each time frame in both fluorescence channels (contour of the particle in cyan). Scale bar, 5 *μ*m. (Bottom) Protocol followed to estimate the protein coverage on the tubes or buds over time, in order to correct for the geometry of lipid structures. **b**, Binding curves of the protein binding to several buds (left) or tubes (right) at 0.25 *μ*M (green colours) or 0.35 *μ*M (orange colours) bulk protein concentration. The protocol described in a and b has been used, enabling to plot the protein coverage on tubes or buds over time. Tube elongation from buds (left) starts when the plot ends, tube pearling (right) starts when the plot ends. **c**, Bud and tube reshaping upon protein adsorption and corresponding protein average coverage on the membrane protrusions.

## Supplementary Videos

**Suppl. Video 1.** Time sequence of the patterned Supported Lipid Bilayer (pSLB) slowly destretching from 5 % strain to 0 %, leading to the formation of lipid tubes. Fluorescence images show the pSLB membrane marker. The hexagonal region drawn in green shows the initial size of the stretched pattern. Note that some defocused frames are kept for the sake of timelapse understanding; as destretch occurs the PDMS membrane moves in the vertical direction and refocusing is required.

**Suppl. Video 2.** Short time sequence of the lipid tubes and buds formed through pSLB destretch. Fluorescence images show the pSLB membrane marker. Inset shows a magnification of the region marked with a green square.

**Suppl. Video 3.** Longer time sequence of the lipid tubes and buds formed through pSLB destretch. Note that tubes relax to buds with time. Fluorescence images show the pSLB membrane marker. Inset shows a magnification of the region marked with a green square.

**Suppl. Video 4.** Time sequence of the lipid tubes and buds formed through pSLB destretch before and after incubation with 1 μM fluorescent neutravidin. Fluorescence images show the membrane marker (left) and neutravidin marker (right). Insets show magnifications of the regions marked with a square.

**Suppl. Video 5.** Time sequence of a non-stretched pSLB before and after incubation with 0.25 μM fluorescent amphiphysin. Fluorescence images show the membrane marker (left) and neutravidin marker (right). Insets show magnifications of the regions marked with a square.

**Suppl. Video 6.** Time sequence of a non-stretched pSLB before and after incubation with 5 μM fluorescent amphiphysin. Fluorescence images show the membrane marker (note that to inject the protein at high concentration, the non-fluorescent form was used). Inset shows magnifications of the regions marked with a green square.

**Suppl. Video 7.** Numerical simulation of the reshaping of a lipid bud of initial diameter 850 nm following protein binding from a protein bulk concentration of 0.5 μM. Color is protein coverage (left) and order (right). Membrane tension is fixed to that prior to protein exposure and the volume enclosed between the membrane and the substrate is fixed.

**Suppl. Video 8.** Numerical simulation of the reshaping of a lipid bud of initial diameter 850 nm following protein binding from a protein bulk concentration of 0.5 μM. Color is protein coverage (left) and order (right). Membrane tension is fixed to that prior to protein exposure and the exchange of the volume enclosed by the protrusion is eased by considering a softer substrate interaction.

**Suppl. Video 9.** Time sequence of the reshaping of lipid tubes and buds formed through pSLB destretch, before and after incubation with 0.5 μM Amphiphysin. Fluorescence images show the membrane marker (left) and Amphiphysin marker (right). Insets show magnifications of the regions marked with squares.

**Suppl. Video 10.** Time sequence of the reshaping of lipid tubes and buds formed through pSLB destretch, after incubation with 0.25 μM Amphiphysin. Fluorescence images show the membrane marker (left) and Amphiphysin marker (right). Insets show magnifications of the regions marked with squares.

**Suppl. Video 11.** Numerical simulation of the reshaping of a lipid tube of initial diameter 260 nm following protein binding from a bulk concentration of 0.5 μM. Color is protein coverage (left) and order (right).

**Suppl.Video 12.** Time sequence of the reshaping of lipid tubes and buds formed through pSLB destretch, before and after incubation with 0.25 μM amphiphysin. Fluorescence images show the membrane marker (left) and amphiphysin marker (right). Insets show magnifications of the regions marked with squares.

**Suppl. Video 13.** Time sequence of the reshaping of lipid tubes and buds formed through pSLB destretch, before and after incubation with 0.35 μM amphiphysin. Fluorescence images show the membrane marker (left) and amphiphysin marker (right). Insets show magnifications of the regions marked with squares.

**Suppl. Video 14.** Time sequence of the reshaping of caps formed through pSLB destretch followed by a hypo-osmotic shock, before and after incubation with 1 μM Amphiphysin. Fluorescence images show the membrane marker (left) and Amphiphysin marker (right). Insets show magnifications of the regions marked with squares.

**Suppl. Video 15.** Time sequence of the reshaping of caps formed through pSLB destretch followed by a hypo-osmotic shock, before and after incubation of 3 μM Amphiphysin concentration at the first indicated injection, and 5 μM at the second one. Fluorescence images show the membrane marker (left) and Amphiphysin marker (right). Insets show magnifications of the regions marked with squares.

**Suppl. Video 16.** Time sequence of human dermal fibroblasts co-transfected with GFP-Mem (left) and mCherry-Amphiphysin (right) before, during and after stretching. Insets show magnifications of the regions marked with squares.

## Notes

### Competing Interest Statement

The authors have declared no competing interest.

